# Impact of Inert Crowders on Host-Guest Recognition Process

**DOI:** 10.1101/2022.03.03.482924

**Authors:** Bibhab Bandhu Majumdar, Jagannath Mondal

**Affiliations:** Tata Institute of Fundamental Research, Hyderabad 500046, India

## Abstract

Biological environments typically contain high concentrations (300-400 mg/mL) of different macromolecules at volume fractions as large as 30%-40%. Biomolecular processes, especially ubiquitous recognition phenomena, occurring in such crowded heterogeneous media would differ significantly compared to the dilute buffer solutions. Here we quantify the impact of inert crowders on prototypical host-guest recognition process by explicit-solvent molecular dynamics (MD) simulations in atomic resolution. We demonstrate that the crowders, especially when smaller in size, would facilitate the binding process of guest molecule by decreasing the free energy barrier for binding via excluded volume effect and desolvation of the host receptor. However, the extent of crowder-induced stabilization of host-guest complex is found to be significantly higher when the guest molecule is sterically constricted to approach the host along a centrosymmetric direction, contrary to its unrestricted, freely diffusive movement. A kinetic analysis of the recognition process reveals that the origin of relatively stronger crowder impact, during constricted movement of guest molecule, lies in significantly enhanced residence time of the guest inside the host by crowders, compared to freely diffusive ligand movement. Together our results suggest that the extent of im-pact of crowding on recognition processes would be contingent upon presence or absence of constriction on ligand movement.

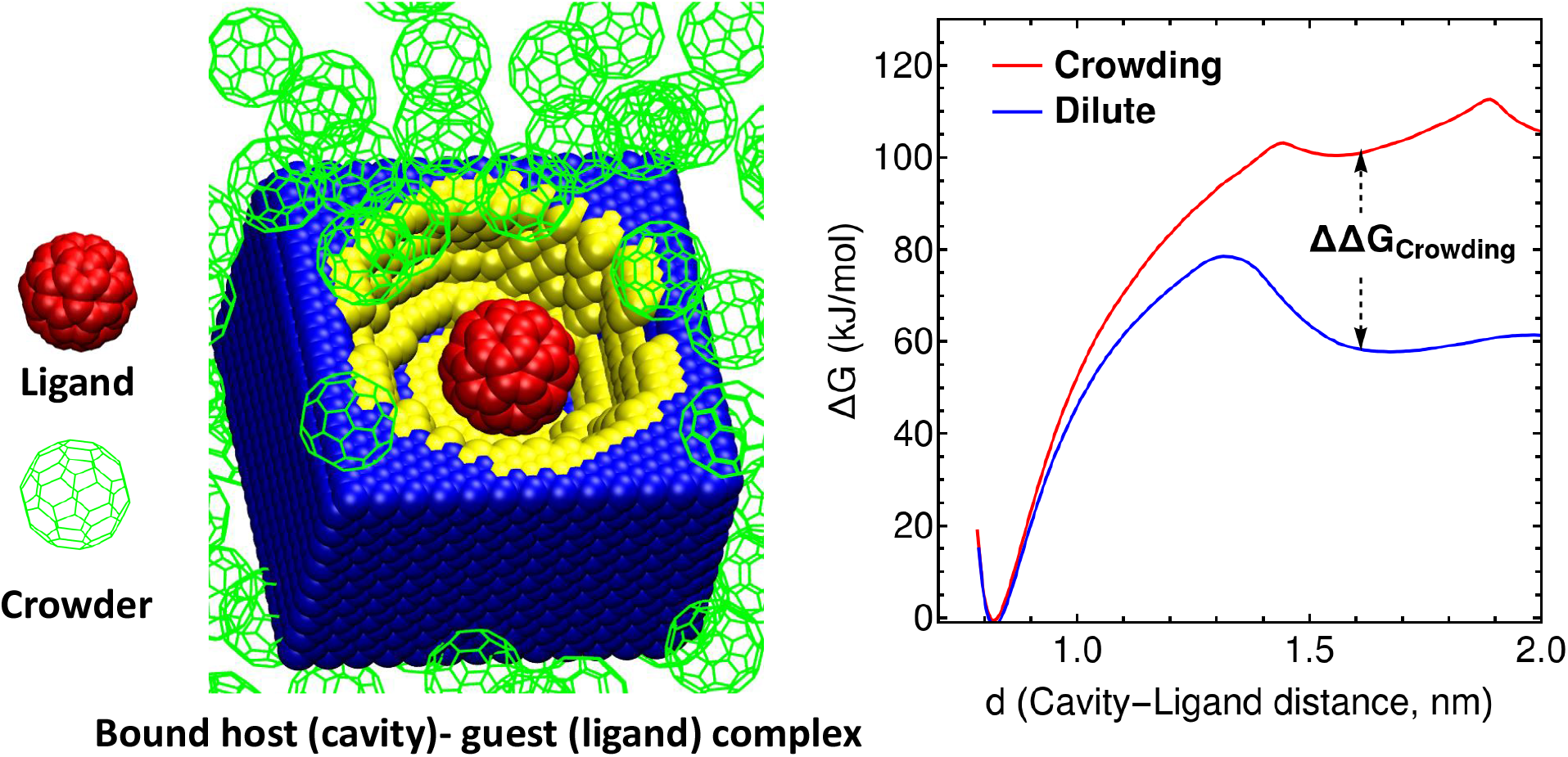
TOC graphic

## Introduction

The term “macromolecular crowding” was coined to underline the impact of volume exclusion by one macromolecule to another macromolecule.^1,2^ The mutual impenetrability of macromolecules can impact various biomolecular processes, such as: biomolecular association, protein ligand binding, biomolecular recognition, protein folding-unfolding equilibria, biomolecular diffusion etc.^3–8^ In this study we are focused on investigating the effect of macro-molecular crowding on host-guest recognition process. Biomolecules in intracellular medium encounter various macromolecules in addition to their specific binding partners. Non-specific interactions between these bystander crowder molecules and proteins are comprised of a multitude of various forces, such as: electrostatic interactions, hydrogen bonding interactions, hydrophobic interactions, van der Waals repulsive interactions. ^7^ Repulsive volume exclusion effect has been established as one of the dominant factors influencing macromolecular crowding phenomena. ^9,10^ Excluded volume effect stabilizes biomolecules with smaller surface-to-volume ratio that effectively facilitates biomolecular association and complex formation.^3,11,12^

Recently few experimental and simulation studies have been reported focusing on crowding effect on protein-ligand binding or biomolecular recognition processes. Ghosh et al. have shown the competitive inhibition effect imposed by addition of Ficoll 70 crowder molecules in maltose-binding-protein solutions using fluorescence titration, ^1^H-^15^N-TROSY NMR spectroscopy and classical MD simulation.^13^ Similarly Luh et al. have shown that ^1^H-^15^N-HMQC well-resolved NMR peaks of WW-domain of pin1 protein disappeared in intracellular media, but regained by adding ovalbumin protein.^14^ Duff et al. have demonstrated the impact of protein and synthetic polymer crowders on enzymatic activity of dihydro folate reductase (DHFR) enzyme based on non-specific crowder-ligand interactions reducing enzymatic activity.^15^ Recently Matic et al. have shown that purely entropic pushing effect or excluded volume effect exerted by crowders (PEG and Ficoll 70) can interfere with ligand accessibility to active sites of lactate dehydrogenase (LDH) enzyme resulting in uncompetitive inhibition.^16^ It has been shown that for hairpin ribozyme, various crowding agents, for example: polyethylene glycol or PEG, dextran, Ficoll 70 etc., proteins, cellular extracts, increase the stability of the most compact state, which is the docked ligand-bound active state of the enzyme. This is achieved exclusively by the enhancement of the docking rate constant.^17^ Macromolecular crowding can also affect the enzyme activities in different systems by favoring the active dimeric form of enzymes.^18–20^ The enzymatic activities of DNA-binding proteins are enhanced significantly in presence of crowding.^21,22^ A recent study by Kasahara et al. has shown that the major binding pathway of PP1-inhibitor to src-Kinase protein is altered in presence of bovine serum albumin or BSA protein crowders.^23^ This has been attributed to non-specific attractive interactions between the crowder and target protein as well as trapping of ligands on the crowder protein surface. This is also in line with a recent study of Micoplasma genitalium cytoplasm modelling, where charged or large metabolites were found to be trapped on protein surface while hydrophobic substrates were found to remain mostly solvated.^24^ These studies indicate that crowders can affect ligand binding processes in various ways by interacting with the ligand or receptor molecule. The nature of this interaction can be purely repulsive or dominant excluded volume effect and weakly attractive. For a purely repulsive interaction regime, crowders are typically shown to facilitate biomolecular associations by increasing effective concentrations of substrates. ^23,25^ Zhou and Minton have employed scaled particle theory (SPT) for hard-sphere particles to obtain quantitative analytic expressions describing the impact of crowding on ligand-binding process.^3,6^

In this study we have explored the aspect of repulsive excluded volume effect exerted by crowder molecules on a prototypical host-guest recognition process by performing classical molecular dynamics (MD) simulations in explicit water with full atomistic resolution. A previously well studied system containing an immobile host receptor site with a semi-ellipsoidal cavity and a sterically constrained C60-Fullerene guest ligand weakly interacting with water, was chosen to represent the ligand binding process. ^26,27^ Over the last one decade, this system has remained a popular model system for investigating the ligand-binding kinetics in this prototypical system using full atomistic classical MD simulations and multiple ensemble based approaches ^26–28^ and well-known for displaying solvent-mediated interactions. We have employed the same system and introduced inert crowders to show the impact of crowding on the thermodynamics and kinetics of ligand binding-unbinding process via classical all-atom unbiased MD simulations and free energy simulations. Our observation shows that the crowders would facilitate the binding process of guest molecule by decreasing the free energy barrier for binding via excluded volume effect. The crowders were also found to facilitate the desolvation process of the host receptor via occlusion of the receptor. The impact decreases with increased size of the crowders. However, the extent of crowder-induced stabilization of host-guest complex depends on how the guest molecule is approaching the host: the impact of crowders is found to be significantly higher when the guest molecule is sterically restricted to approach the host along a centrosymmetric direction, contrary to its unrestricted movement. A kinetic analysis of the recognition process reveals that the origin of such steric-controlled crowding effect lies in significantly differential enhancement of the residence time of the guest inside the host by crowders. In contrast, the binding time or on-rate constants have much less effect. Simulation results were further compared with predictions from widely used analytical model for macromolecular crowding: scaled particle theory.^29^

In the next sections the paper is organized as follows: a brief discussion on description of simulation systems and methods employed, results obtained and conclusion of this study. This study can provide useful insights when studying biomolecular recognition in vivo environments where free movement of ligand molecules are sterically hindered in presence of high concentrations of bystander molecules.

## Method and Model

### Model system

We have selected a previously well-studied prototypical cavity-ligand system consisting of a semi-ellipsoidal cavity cut from a rectangular receptor site and C60-fullerene ligand in presence of explicit water molecules.^26,27^ In Fig. 1 we have shown different components of the model system. The receptor molecule consists of two different types of atoms: wall atoms or CW (blue) and cavity atoms or pocket atoms (CP, yellow). The ligand or guest is shown as a red spherical object with carbon atoms represented by van der Waals spheres.

**Figure 1:**
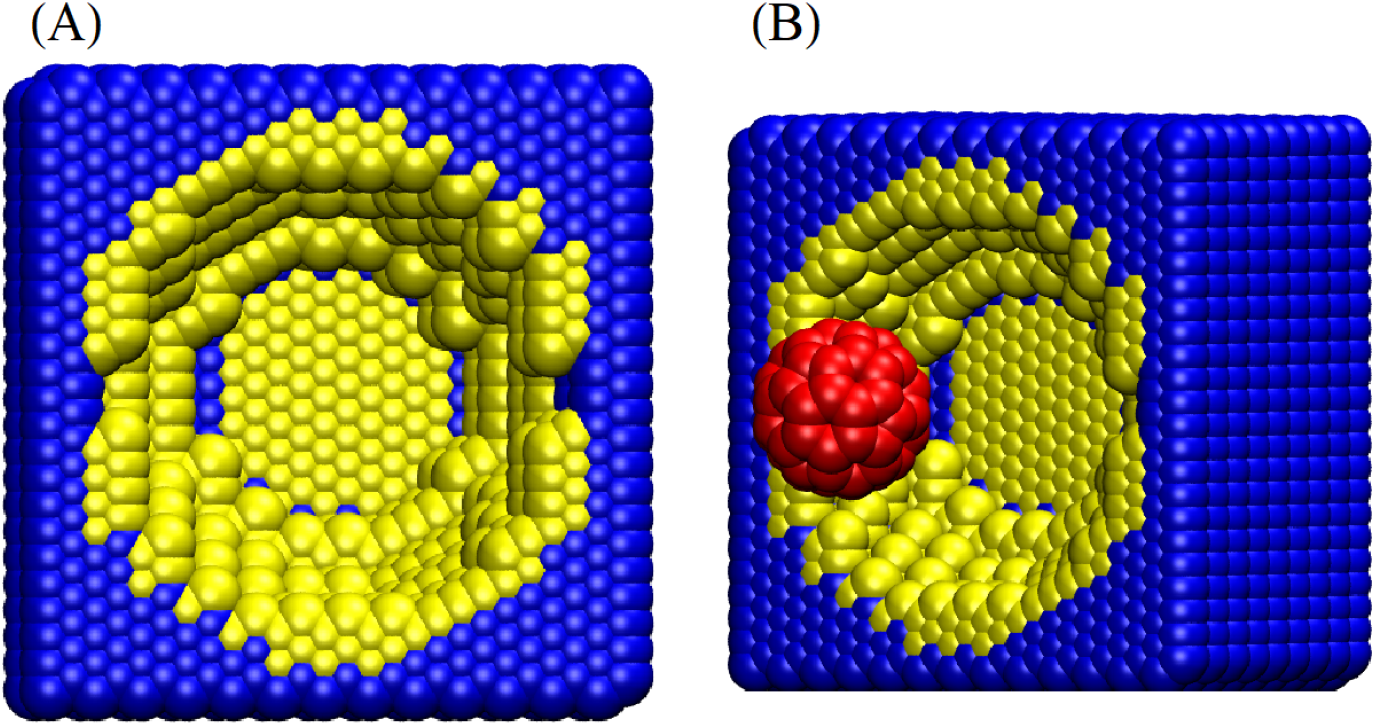
(A) and (B) shows structure of the receptor in absence and presence of a C60-fullerene ligand (red, CX-atom) separated at a certain distance from the receptor. Panel (A) shows structure of the receptor: yellow atoms (CP-atom) constitute the semi-ellipsoidal pocket and blue atoms (CW-atom) constitute rest of the receptor molecule. All atoms are represented as van der Waals spheres (van der Waals radii of atoms are: *r*_*CP*_ = *r*_*CW*_ = 0.4152 nm, *r*_*CX*_ = 0.35 nm)

We have prepared two different sets of simulation systems, namely: dilute (without crowders) and crowded (with crowders). In majority of the simulations described here, an inert crowder of the same geometry as ligand (i.e. same size of C60) was employed. To investigate the effect of crowder-size on host-guest complexation process, we have also performed a set of control simulations using larger sized crowder equivalent to the topology of C240-fullerenes. In Fig. 2 we have shown three representative simulation systems, namely: dilute (in absence of crowders) and crowded-in presence of C60 and C240-crowders respectively, in presence of explicit water (shown as small red spheres). The C60-fullerene structure was obtained from earlier studies of the prototypical cavity-ligand system.^27^ The C240-fullerene structure was obtained from an online repository:https://nanotube.msu.edu/fullerene/fullerene-isomers.html. of fullerene structures. Thus obtained C240 structure was kept in a cubic box with box vector = 4 nm and then solvated with water molecules. This was followed by an energy minimization with steepest descents. The energy minimized C240-fullerene structure was later used as C240-fullerene crowder in our simulations.

**Figure 2:**
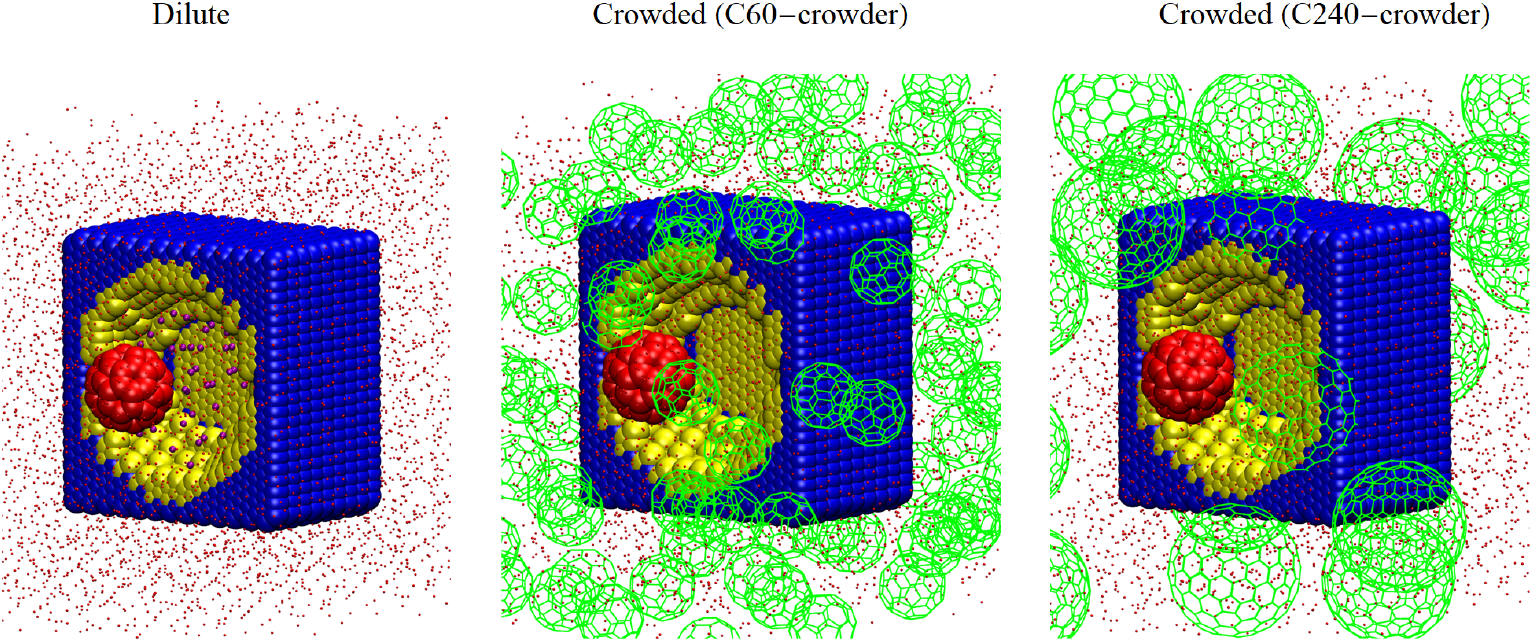
Three different simulation systems are shown: Dilute (extreme left) or in absence of crowders, crowded-in presence of C60-fullerene crowders) and crowded-in presence of C240-fullerene crowders. Crowders are shown as green spheres; water molecules are shown as small red spheres and slightly larger purple spheres in Dilute-system refer to the pocket or cavity water (water molecules trapped inside the cavity); Cavity and ligand atoms are shown in similar representations as described in Fig. 1

Bonding and non-bonding interaction energy parameters for the interactions between receptor, C60-fullerene ligand and water were taken from an earlier study on similar system ^26^ Potential well depth parameters for the semi-ellipsoidal cavity atoms (CP-atom) and wall atoms (CW-atoms) were set to 0.008 and 0.0024 kJ mol^−1^ to obtain “intermediate-attractive” interactions between the receptor and the ligand. ^27^ The interactions between water and both cavity and ligand atoms are weak showing hydrophobic nature of the system. Attractive pairwise LJ interactions between crowder-crowder, crowder-ligand and crowder-receptor were excluded to keep the interactions between the crowder and any other component of the simulation system (except water) purely repulsive and prevent self-aggregation among the crowders. Thus crowder-crowder, crowder-ligand and crowder-receptor interactions were kept purely repulsive except for the crowder-water interactions which were set to typical 12-6 LJ interactions. Intramolecular interactions between ligand atoms and receptor atoms were excluded. Motions of the host receptor in all three directions were discarded to keep the receptor fixed at same position during the course of the simulations. TIP4P water model^30^ was used to describe water molecules in the system. Number of water molecules in the simulation were ranging from 4000 to 6000 for crowding and dilute solutions respectively.

## Simulation setup and protocols

We have simulated multiple scenarios in presence and absence of crowding. We have further categorised the systems based on ligand degrees of freedom. We have prepared two sets of systems where: (1) ligand movement is restricted and ligand can move only along centrosymmetric approach (*z*-direction) and (2) ligand movement is unrestricted and ligand can freely diffuse in any direction. These two situations are described by Fig. 3 for a prototypical ligand binding-unbinding process.

**Figure 3:**
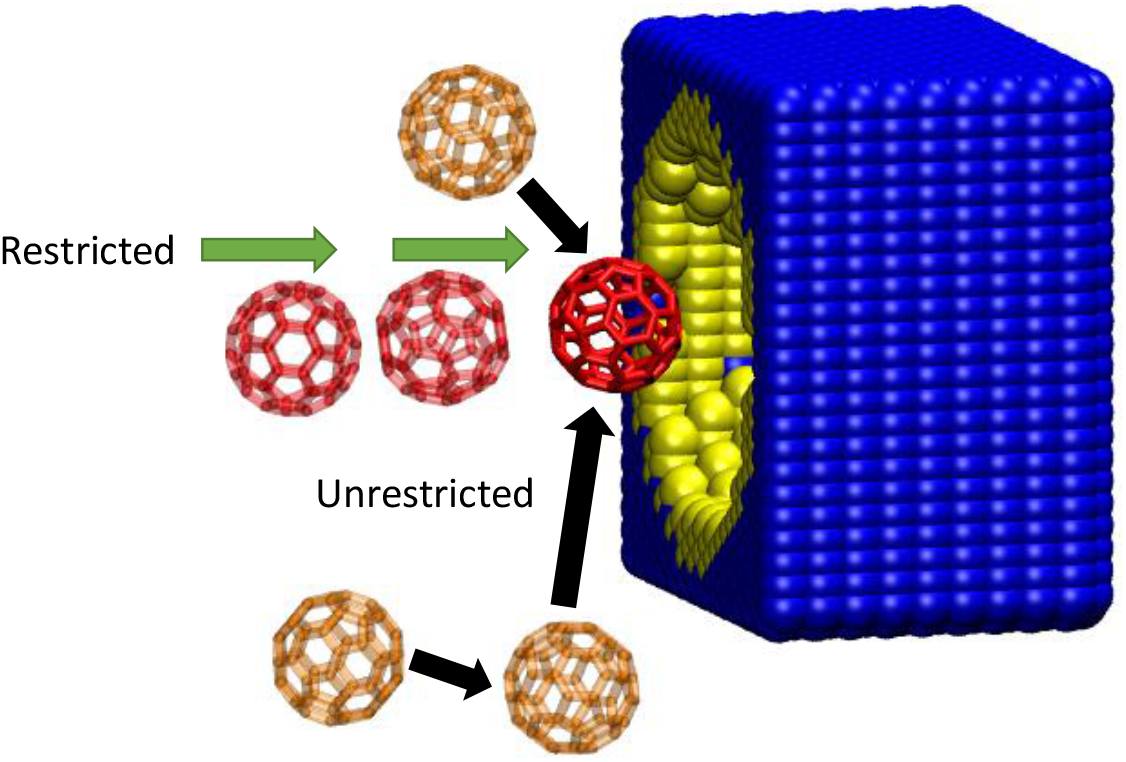
Two sets of systems based on ligand degrees of freedom are shown: solid red-sphere refers to the initial position of the ligand, red transparent spheres along the green arrows refer to the restricted ligand movement and orange transparent spheres along the black arrows refer to the unrestricted ligand movement.

In the following, we will describe the protocols employed to simulate systems for restricted and unrestricted ligand movement.

### Restricted movement of the ligand

Here we have chosen cavity-ligand systems as described in the previous section in presence and absence of macromolecular crowders (C60 and C240-fullerenes). In these systems the ligand was restricted to move only along the z-direction during the course of the simulations.

These model systems represent situations where a ligand is sterically restricted to move along one direction, e.g. in case of host-guest complex such as cucurbituril-adamantane,^31^ ion-channels, membrane proteins etc. The starting configurations were prepared by putting the cavity-ligand system in a simulation box with box vectors 5 nm, 5 nm and 8 nm corresponding to a box volume of 200 nm^3^. The distance between centers of geometry between the receptor and the ligand was chosen as the preferred reaction coordinate to describe the ligand binding-unbinding process. In order to investigate the impact of crowding with respect to dilute solution, multiple crowding simulations at increasing crowding concentrations and a control simulation in absence of crowder i.e. in dilute solution were performed. Crowding simulations were performed for different volume fractions of the crowder (C60 or C240 fullerene-sized crowders). The van der Waals volumes of C60 and C240 fullerenes: 0.549 nm^3^ and 2.6854 nm^3^, respectively, were obtained from a detailed analytical study of van der Waals volumes and surface areas for different fullerene molecules by Adams et al.^32^ Initial configurations for crowding simulations were prepared by inserting different number crowder molecules corresponding to different crowder volume fractions as described in Table. 1.

**Table 1:** Crowder volume fractions (*ϕ*_*Crowder*_) and number of crowder molecules

| $\phi_{Crowder}$ | $N(C60)$ | $N(C240)$ | box-volume (nm<sup>3</sup>) |
| --- | --- | --- | --- |
| 0.05 | 18 | 4 | 200 |
| 0.10 | 37 | 8 | 200 |
| 0.15 | 55 | 11 | 200 |
| 0.20 | 73 | 15 | 200 |
| 0.25 | 91 | 19 | 200 |

First, we have performed 25 independent unbiased MD simulations at C60-Fullerene crowding volume fractions of 0.25 and in dilute solution (*ϕ*_*Crowder*_ = 0.0). Each unbiased MD simulation was performed by running a steepest descent energy minimization followed by a 25 ns MD simulation in canonical ensemble. Temperature of the system was controlled using velocity rescaling thermostat with a reference temperature of 300 K and time constant of 1 ps.

To quantify the free energetics of host-guest recognition process, we have also performed umbrella sampling^33^ simulations in presence and absence of macromolecular crowding. For this purpose we have chosen C60 and C240-fullerene molecules as crowders. Furthermore we have prepared configurations for umbrella sampling MD simulations. Configurations with different values of the reaction coordinate (receptor-ligand distance) were obtained by placing the ligand fullerene at various distances ranging from 0.76-2.01 nm with 0.05 nm interval from the receptor. This was followed by insertion of the crowders and solvation of the individual configurations. In total we generate 26-starting configurations or umbrella windows for umbrella sampling simulations in presence and absence of crowding. Each configuration was energy minimized with steepest descents followed by a 2 ns and a 10 ns umbrella sampling simulations performed in isothermal-isobaric and canonical ensembles respectively. Temperature of the systems were controlled using velocity rescaling thermostat with a reference temperature of 300 K and time constant of 1 ps. All simulations were performed using the GROMACS-2018 software package. ^34^ The force constant for harmonic restraint on the reaction coordinates ranged between 1000 to 6000 kJ mol^−1^ nm^−2^ and were optimised by verifying sufficient overlap of probability distributions between adjacent windows. The resulting free energy profiles were obtained by reweighing the distributions using a one dimensional weighted histogram analysis method (WHAM) code implemented in the GROMACS software package.^35^

### Unrestricted movement of the ligand

In order to explore impact of crowding on ligand binding process involving a freely diffusive C60-ligand, we have performed multiple biased and unbiased simulations with ligand free to move in any direction.

A total of twenty independent unbiased simulations of 25 ns in explicit water molecules with full atomistic resolution were run for multiple values of the reaction-coordinate in presence and absence of C60-fullerene crowders. In these simulations, the ligand was placed at specific distances from the cavity or placed randomly inside the simulation box. Simulation protocol similar to unbiased simulations of restricted ligand movement case were followed for the unbiased simulations involving unrestricted ligand movement. We have chosen two reaction coordinates to describe the ligand-binding process for these simulations. These are namely, the *z*-distance (distance in the z-direction) and 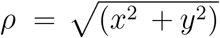 (distance in the xy-plane), where *x, y*, and *z* are the respective components of receptor-ligand distances.

Furthermore we have performed multiple biased two-dimensional umbrella sampling simulations for dilute solution and C60-crowder volume fractions of 0.15, 0.2 and 0.25. Initial configurations were generated by placing the ligand at multiple *z* and *ρ* values ranging from 0.81-2.01 nm and 0-2.0 nm respectively. A *ρ* value of zero refers to the centrosymmetric approach of the ligand, whereas any non-zero value for *ρ* will refer to deviation from the centrosymmetric approach. A total of 156 configurations were generated for the 2D-umbrella sampling simulation. Harmonic restraint potential settings were employed as described in Ref. 26. A two-dimensional harmonic restraint potential with force constants of 4500 kJ mol^−1^ nm^−2^ were used. After an initial energy minimization, a 2 ns umbrella sampling in isothermal-isobaric ensemble was followed by a 2 ns umbrella sampling simulation in canonical ensemble for each configuration. The 2D-free energy surface as a function of the reaction coordinates: *z* and *ρ* was obtained using a 2D-WHAM code.^36^

### Method to calculate rate constants for host-guest complex formation

In order to investigate the impact of macromolecular crowding on the host-guest recognition kinetics, we have calculated the binding (*k*_*on*_) and unbinding rate constants (*k*_*off*_) as described by the process: 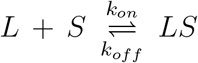. Here *L, S* and *LS* refer to the guest ligand, the host receptor molecule and the receptor-ligand complex, respectively. We have estimated the values of the on and off-rate constants using the following equation:

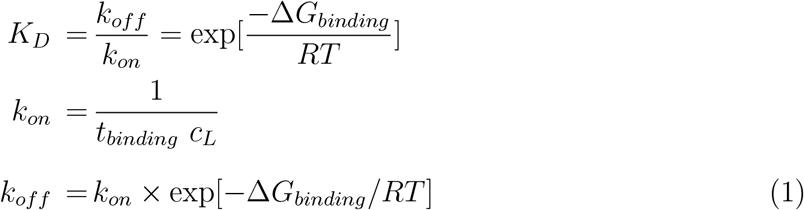

In Eqn. 1, *K*_*D*_ refers to the dissociation equilibrium constant and Δ*G*_*binding*_ (kJ mol^−1^) is the free energy of binding obtained from the umbrella sampling simulations and *c*_*L*_ and *t*_*binding*_ refer to the molar concentration of the ligand and the binding time or the time taken by the ligand to bind to the host receptor, respectively. The binding times are estimated from the time-profile of the reaction coordinate(s) obtained from the unbiased MD simulation.

We have calculated the values of *k*_*on*_ and *k*_*off*_ in dilute (*ϕ*_*Crowder*_ = 0.0) as well as in crowded solutions (C60-fullerene crowder, *ϕ*_*Crowder*_ = 0.25), for both sterically restricted as well as freely diffusing ligand. It is noteworthy, that in dilute solution for sterically restricted movement of the ligand, we have not observed any binding events from multiple independent unbiased MD simulations. This can be attributed to the limited unbiased MD simulation time lengths used for the purpose of this study. Ahalwat et al. have estimated the mean first passage time or binding time for sterically restricted ligand moving along the z-direction in dilute solution in this prototypical host-guest system using a combination of unbiased MD simulations and markov state modeling approach.^28^ We have used the binding time reported in Ref. 28 for calculating *k*_*on*_ and *k*_*off*_ in dilute solution for restricted ligand movement.

### Scaled particle theory(SPT) of host-guest complexation

We compared our simulation results of crowding effect on host-guest recognition process with a popular theory namely scaled particle theory (SPT). Since its inception, ^29^ SPT has been successfully employed to describe the impact of macromolecular crowding on biomolecular processes.^3,6,37,38^ SPT of fluids was introduced by Reiss et al. to study thermodynamical properties of fluids considering them as hard-spherical particles. ^39^ SPT provides exact quantitative analytic estimates for free energy change of inserting an inert, hard test particle in terms of the work required to create a cavity of the size of the test particle in a solution of same or other macromolecules present in arbitrary concentrations.^3,6^ For a thermodynamic cycle describing host-guest complex formation: binding of guest ligand molecule (L) to an immobile host receptor site (S): *L* + *S* ⇌ *LS*, the free energy of binding in presence and absence can be written as:^3,6^

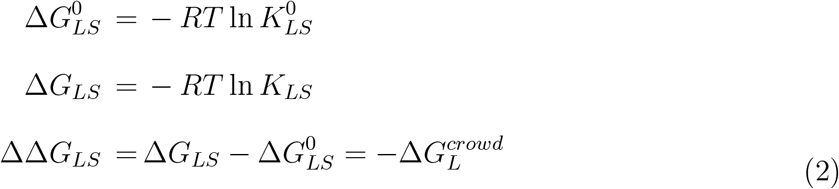

Where ΔΔ*G*_*LS*_ refers to the change in free energy of binding due to macromolecular crowding, 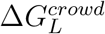 refers to the standard free energy of transferring a guest ligand molecule to a *LS* solution of hard-spherical crowders occupying a volume fraction of: (*ϕ*_*Crowder*_) and 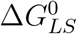 represents standard free energy of binding or host-guest complex formation in dilute solution and in presence of crowders respectively. In Eqn. 2, *R* is the universal gas constant, *T* is the absolute temperature, 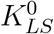 and *K*_*LS*_-are the corresponding equilibrium constants in dilute and crowded solution respectively. SPT provides an expression for estimating the free energy of creating such cavity and is particularly useful in quantitative estimation of macromolecular crowding effects in biomolecular processes.

According to the SPT, the free energy of transferring a hard spherocylindrical guest ligand particle 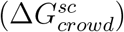 into a solution of hard spherical crowder molecules occupying a volume fraction *ϕ*_*Crowder*_, are described as:^3^

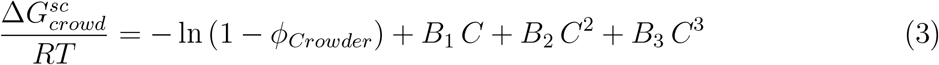

Where *C, B*_1_, *B*_2_ and *B*_3_ can be written as:^3^

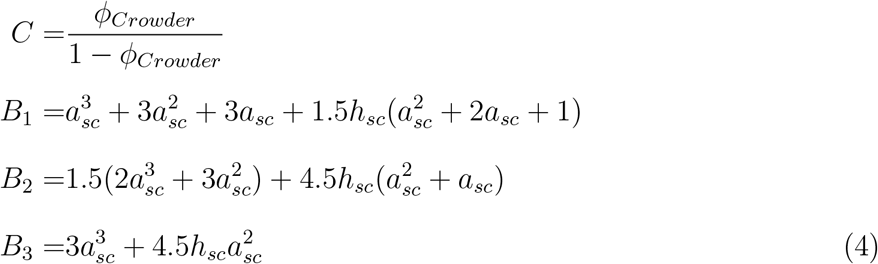

In Eqn. 4, *a*_*sc*_ and *h*_*sc*_ refer to the ratio of radii of the guest ligand and crowder molecule and the length of the guest ligand, respectively. Free energy of transferring a spherical ligand molecule can be obtained by putting *h*_*sc*_ = 0 in Eqn. 4.

We have compared our simulation results with corresponding SPT predictions. Towards this end, from free energy simulations we estimated 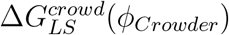 in crowded solution and Δ*G*_*LS*_(0) in dilute solution and got an estimate of ΔΔ*G*_*LS*_. For a detailed description of SPT predictions of crowding on ligand binding process readers are advised to refer to Ref. 3.

## Results and Discussion

In this section we will discuss effect of macromolecular crowding on the ligand binding process. For this purpose we have chosen the distance between the center of geometries of the ligand and entire receptor molecule as reaction coordinate. In the following sections we have discussed the key findings obtained from MD simulations of the cavity-ligand system in presence and absence of macromolecular crowding for both restricted and unrestricted ligand movement.

### Impact of crowders in case of restricted movement of guest towards host

In this section we have shown the impact of crowding on ligand binding when the guest is restricted to move along a centrosymmetric direction towards the host. At first, we have computed the time profiles of the reaction coordinate obtained from multiple independent unbiased simulation trajectories in presence of crowders at a range of crowder volume fractions (*ϕ*_*Crowder*_) and compared the same with that of dilute, uncrowded solution (figure 4).

**Figure 4:**
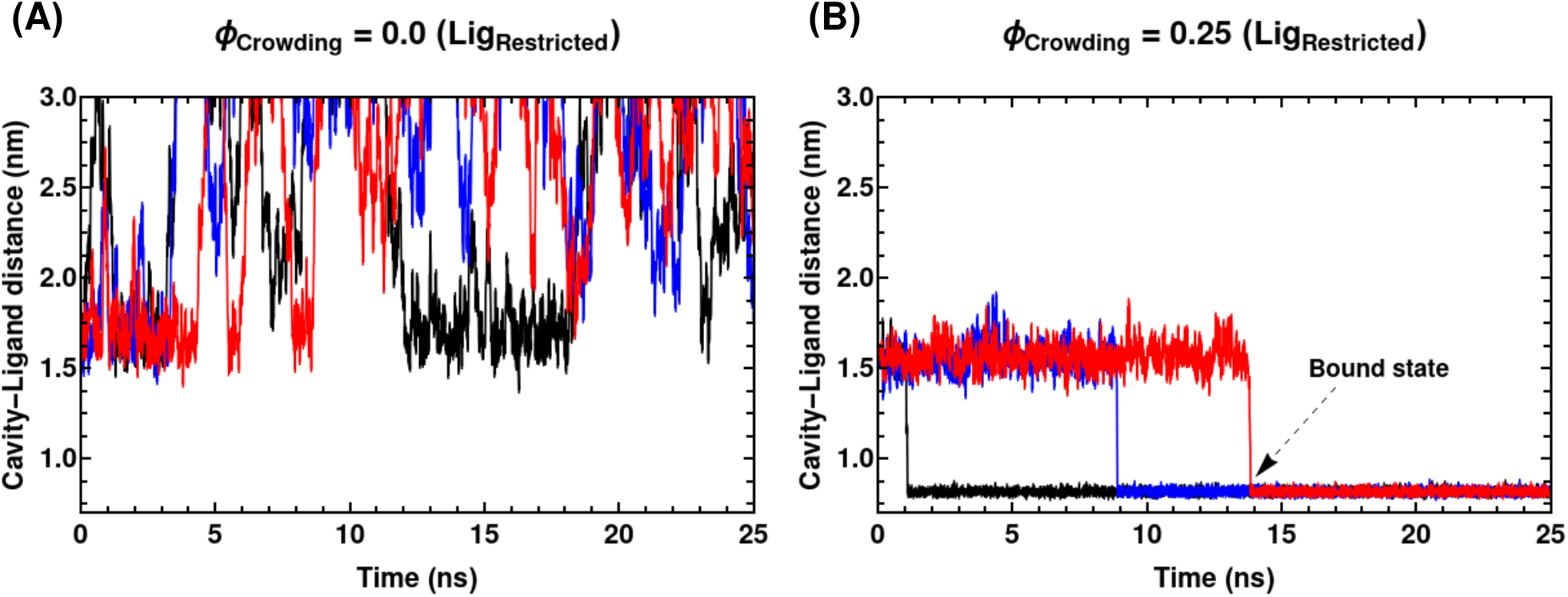
Time evolution of cavity-ligand center-to-center distances in presence (*ϕ*_*Crowder*_ = 0.25) and absence (*ϕ*_*Crowder*_ = 0.0) of crowding obtained from unbiased MD simulations. C60-Fullerene is used as crowders in these simulations. Multiple independent trajectories of the same system are shown together (blue, black and red lines). The dashed arrows refer to the ligand bound state.

In fig. 4 time profiles of the cavity-ligand distances are shown in presence and absence of crowding. In absence of these inert crowders, as reflected by the distance profiles in fig. 4 A, the ligand is not able to reach to the cavity within the similar simulation time. Inability of the guest ligand to bind to the host receptor within limited simulation time length indicates the presence of impending high energy barrier that the guest molecule needs to circumvent prior to binding at the designated cavity. In fact, recent investigation based on a Markov state modelling (MSM) of binding process of the same system^28^ had predicted significantly long binding time (285 nanosecond) for this ligand. However, it is noteworthy that with increase in *ϕ*_*Crowder*_, the average time for onset of binding gradually decreases. As can be seen from Fig. 4B, the distance-profiles reach a plateau region at approximately 0.8-0.9 nm at different simulation times in presence of crowders at *ϕ*_*Crowder*_ = 0.25. We will consider this distance range of 0.8-0.9 nm as the distance where ligand is completely bound to the receptor. This indicates that with increase in crowding intensity the thermodynamic barrier to binding may gradually decrease. A detailed analysis of binding and unbinding rate constants is reserved for the later part of the article.

In order to quantitatively compare the free-energetics of ligand-binding phenomena across various crowded solution, we have further compared the free energy profiles, Δ*G* (kJ mol^−1^), obtained from umbrella sampling simulations.

In Fig. 5 A), free energy of ligand-binding are compared across a range of crowded solution of various concentration. We have also compared these free energy profiles with those in presence of solution constituted of larger (C240) fullerene crowder (keeping the ligand same) (Fig. 5 B)). In Fig. 6, we have shown the free energies associated in the host-guest recognition process: free energy of binding (Δ*G*_*binding*_), barrier towards binding 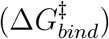 and the barrier towards unbinding 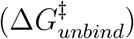, respectively for both crowded and dilute solution. In Table. 2 we have shown the free energies in presence of both C60 and C240-Fullerene crowders.

**Table 2:**
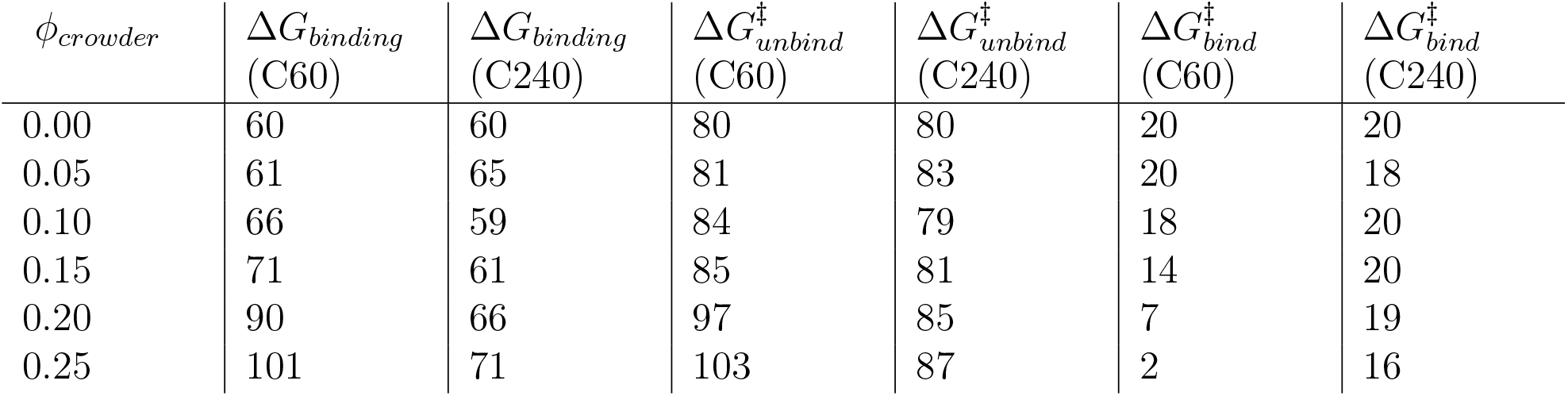
Crowder volume fractions (*ϕ*_*crowder*_), free energy of binding (Δ*G*_*binding*_), barrier to unbinding 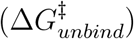; and barrier to binding 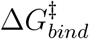 are shown for C60 and C240-fullerene crowders; Free energies are in units of kJ mol^−1^

**Figure 5:**
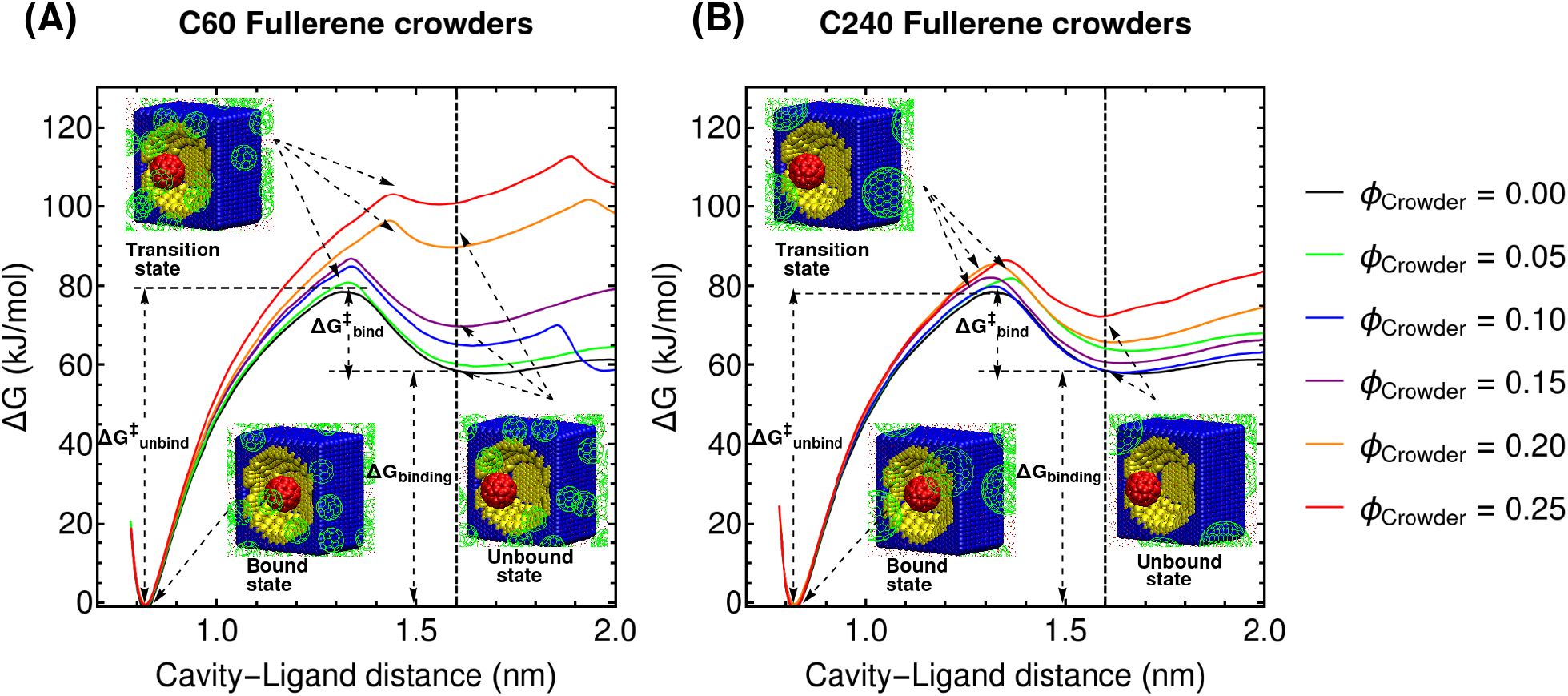
Free energy profiles (Δ*G* (kJ mol^−1^)) for ligand-binding process in presence of C60 and C240-fullerne crowders at various crowder volume fraction. In both cases the Δ*G* value in dilute solution or in absence of any crowder (black curve, *ϕ* = 0.0) is shown for comparison. Three representative crowding simulation snapshots are shown to depict the bound, transition and unbound states in left and right panels. The dashed line at ligand-cavity separation distance of 1.61 nm, refers to the unbound state in different simulations; 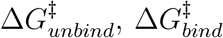 and Δ*G*_*binding*_: free energies associated with the ligand binding processes for *ϕ*_*Crowder*_ = 0 are shown (details explained in text).

**Figure 6:**
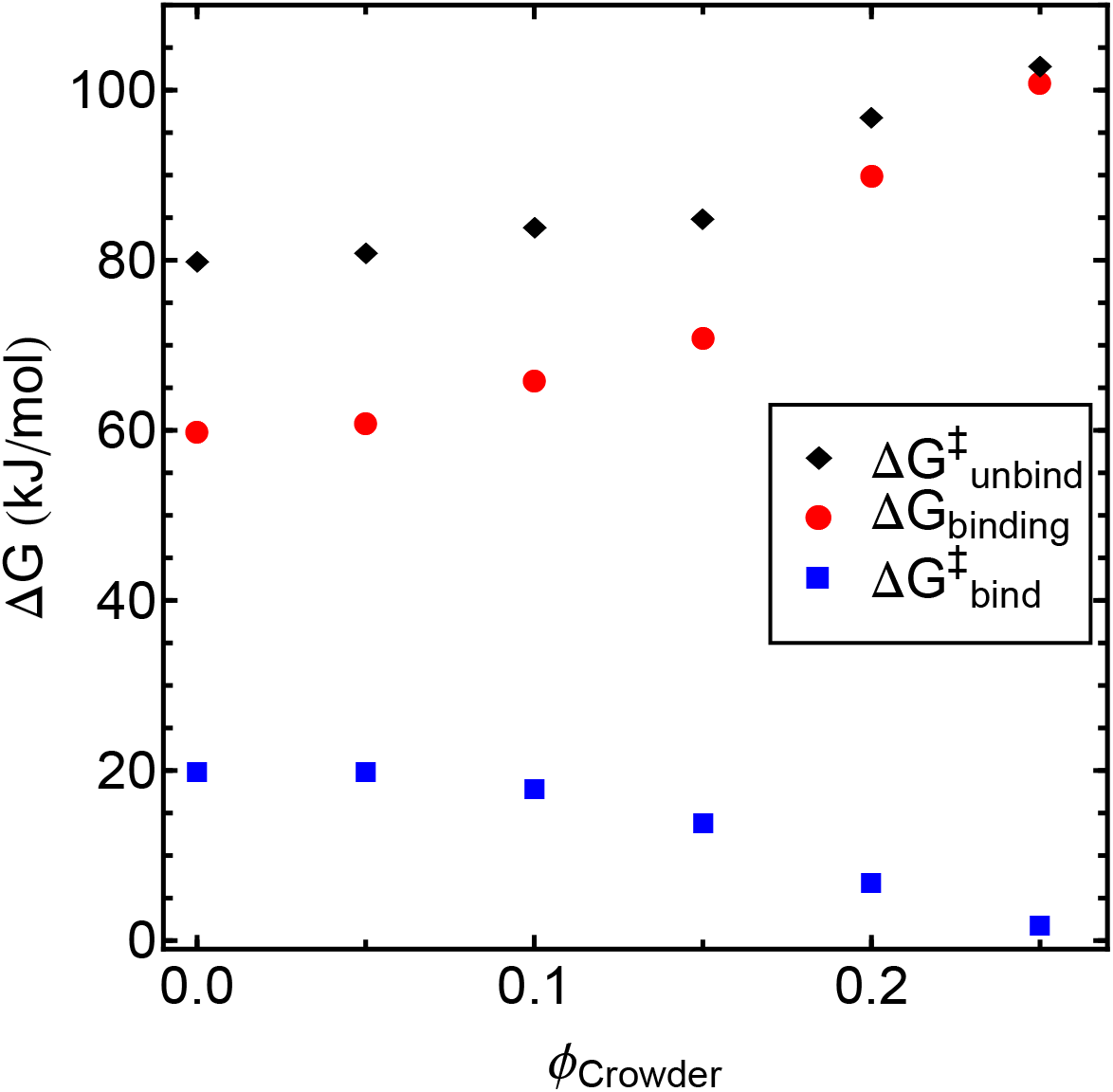
Free energies (Δ*G*_*binding*_, 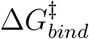 and 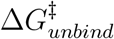 associated with the host-guest recognition process in presence and absence of C60-Fullerene crowders for various crowder volume fractions (*ϕ*_*Crowder*_)

Fig. 5 and Fig. 6 show that with increasing crowding intensity i.e. with increasing *ϕ*_*Crowder*_, the free energy barrier for unbinding 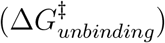 and binding 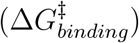 gradually increases and decreases respectively, resulting in overall increase in free energy of binding (Δ*G*_*binding*_). Here 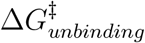 and 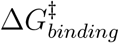 are defined as the barrier to overcome for transitioning from the bound to the transition state and from the unbound to the transition state, respectively. This can be justified as a consequence of excluded volume effect where ligand-receptor association is entropically favorable to avoid energetically unfavorable contacts with the crowders. In order to minimize crowder contacts, ligand prefers bound states over unbound states with increase in crowding concentration.

However, It can be noted from Fig. 5, a very mild stabilization of unbound states starts to appear at very large receptor-ligand separation (*>* 1.9 nm). In fact, it is noteworthy that increasing crowder concentration arbitrarily will not always ensure facilitated binding process. In order to verify impact of very high crowding concentrations on binding process, we have performed additional umbrella sampling simulations for a relatively high crowder concentration (*ϕ*_*Crowder*_ = 0.31) and the corresponding free energy profiles are shown in the supplementary information (Fig.S1), which predicts stabilisation of a relatively unbound state at larger host-guest separation. We have further calculated average number of crowder-ligand contacts as function of the reaction-coordinate for various volume fractions of C60-sized crowder. For low concentration of crowders (*ϕ*_*Crowder*_ = 0.05 − 0.20) the average number of contacts are smaller in magnitude and does not show large fluctuations over the entire range of reaction-coordinate. However, larger crowder-ligand contacts are observed for *ϕ*_*Crowder*_ *>* 0.2, significantly larger number of contacts are observed for ∼31 % volume fractions at larger ligand-receptor separations. The observation evidently depicts that when ligand is approaching the receptor from a very large distance, very large crowder concentration may lead to enhanced crowder-ligand contact events. This eventually occlude the ligand to bind to the receptor. This implies that increasing crowder concentration arbitrarily, will not always result in facilitating the ligand-binding process. In presence of very high crowding concentration, eventually reduced ligand diffusion can decelerate binding compared to dilute solution scenario. Crowding can impact a biomolecular process by: (1) reducing the energy of the transition-state (reaction or activation controlled) and (2) reducing the diffusivity of reactants involved (diffusion controlled). ^3,40,41^ Zhou et al. predicted macromolecular crowders to be most effective when the process under study is activation controlled (slow associations) rather than diffusion-controlled (fast associations). ^3^ Impact of a crowder on a biomolecular process depends on trade-off between these two effects. ^42^

Crowder size can also have significant impact on ligand-binding processes. Left and right panels of Fig. 5 show the different intensity with which two different sized crowders impact the free energetics of the ligand-binding process. Increase in barrier to unbinding and Δ*G*_*binding*_ with *ϕ*_*Crowder*_ is much more pronounced in presence of C60 compared to C240. This trend may seem counterintuitive owing to the fact that C240 is much larger in size compared to C60. The number of crowder molecules will decrease with increase in crowder size for a specific volume fraction. The number density of crowders will be much larger for C60 than C240-crowders in order to attain a specific value of *ϕ*_*Crowder*_. Thus, the ligand-binding process will be more strongly perturbed in presence of a smaller crowder i.e. C60 compared to a larger crowder, C240.

In Fig. 7 we have shown the probability densities of LJ interaction energies between crowder-crowder and crowder-ligand in presence of both C60 and C240 crowders at a volume fraction of 0.25. Panel A of Fig. 7 shows that crowder-crowder repulsive interactions are significantly more stronger in presence of C60 compared to C240 crowders. This effectively gives rise to relatively stronger crowding intensity for C60 crowders. Panel (B) of Fig. 7 shows that in addition to crowder-crowder interactions, contacts between crowder and ligand are also significantly less repulsive or less unfavorable in presence of C240 crowders. This in turn allows the ligand to diffuse along the z-direction relatively at more ease in presence of larger crowder, giving rise to smaller barrier to unbinding and smaller free energy of binding for C240 crowders. A balance between crowder size, number density and interaction energies makes smaller crowders much more effective compared to a larger crowder. In addition to this, impact of crowder size can also be argued in terms of preferential interactions between the solute and the crowders.^37,43,44^ It has also been argued that chemically softer repulsion between crowders can actually induce effective attraction between the crowder and the solute molecules due to enhanced depletion between softly repulsive crowders.^45^ We have shown that crowder-ligand interactions are less unfavorable in presence of C240-fullerene crowders compared to C60-fullerene crowders. In an earlier study Sharp has shown that larger crowders effectively decrease solute chemical potential coefficients (curvature, volume and surface coefficients of solute chemical potential). Preferential interactions of the ligand with larger crowders over smaller ones lead to stabilization of unbound configurations. Our results agree with the qualitative predictions of Sharp et al.,^37^ which essentially predicts smaller crowders can perturb the outcome of a biomolecular process relatively more strongly compared to larger crowders at constant crowder volume fraction. The crowder size dependence can also be described in the light of the depletion forces theory. It has been shown that in accordance with the Asakura-Oosawa (AO) model of depletion forces, at constant volume fraction, smaller cosolute or crowders can result in larger net attractive interaction between two solute surfaces.^37,46^ A recent study highlighted the impact of crowder size on RNA tertiary folding using single molecule Föster resonance energy transfer or smFRET experiment in presence of crowders of varying molecular weights, where smaller crowders were shown to increase the protein folding rate and decrease the unfolding rate, significantly compared to larger crowders.^47^

**Figure 7:**
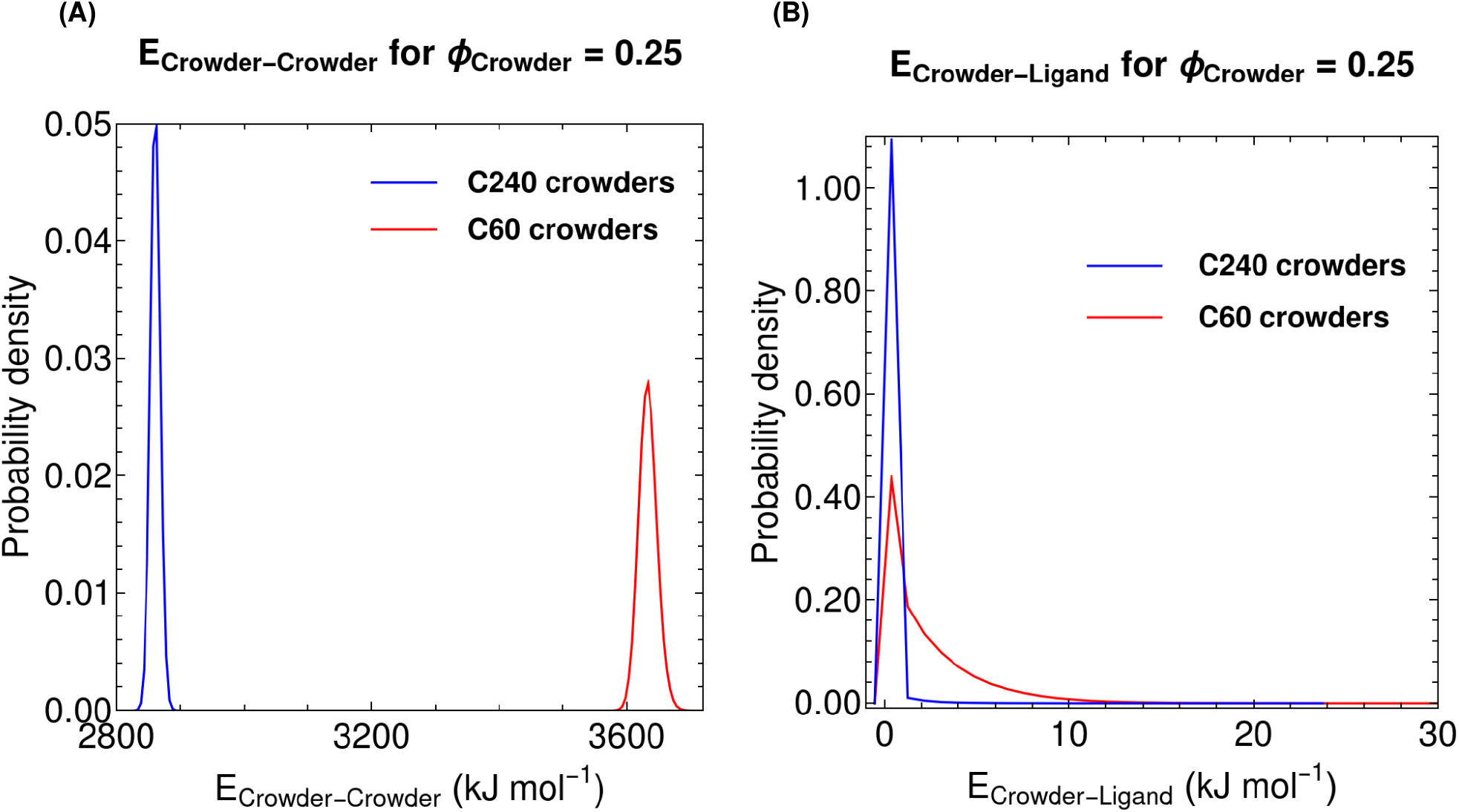
Discrete Probability densities of average crowder-crowder and crowder-ligand interaction energies in presence of C60 and C240-sized crowders at *ϕ*_*Crowder*_ = 0.25.

The impact of crowder size on ligand binding process has also been accounted by the SPT of hard sphere fluids mixtures that predicts larger crowders to have smaller impact on biomolecular processes. ^3^ In later sections we have elaborated on the analytical description of crowding based on the SPT in more detail. Crowder shape is also an important factor influencing biomolecular processes that has not been considered in this study. Largely aspherical crowders can exert a very different impact on a biomolecular processes compared to spherical crowders.^45,48,49^ We compare our simulation results with a well-established analytical model of crowding: scaled particle theory (SPT) of hard-sphere fluid mixtures.

In Fig. 8 we have compared the change in free energy of binding due to macromolecular crowding. We have measured ΔΔ*G* using the following relation from our umbrella sampling simulations:

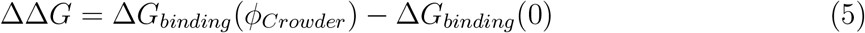

where Δ*G*_*binding*_(*ϕ*_*Crowder*_) refers to the free energy of binding or Δ*G*_*binding*_, which is obtained from umbrella sampling simulations of ligand-binding process in presence of crowder molecules at various volume fractions and Δ*G*_*binding*_(0) is obtained from umbrella sampling simulations in dilute solutions i.e. in absence of crowders. SPT estimate of ΔΔ*G* can be obtained using Eqn. 3 where only dependence on relative sizes of crowders will be changed according to the ratio of the sizes of ligand and crowders where both ligand and crowders are assumed to be spherical. Simulation results for restricted ligand movement for both C60 and C240-crowders qualitatively agree with SPT predictions depicting that in presence of crowding, the ligand-binding equilibrium will be more facilitated compared to dilute solution.

**Figure 8:**
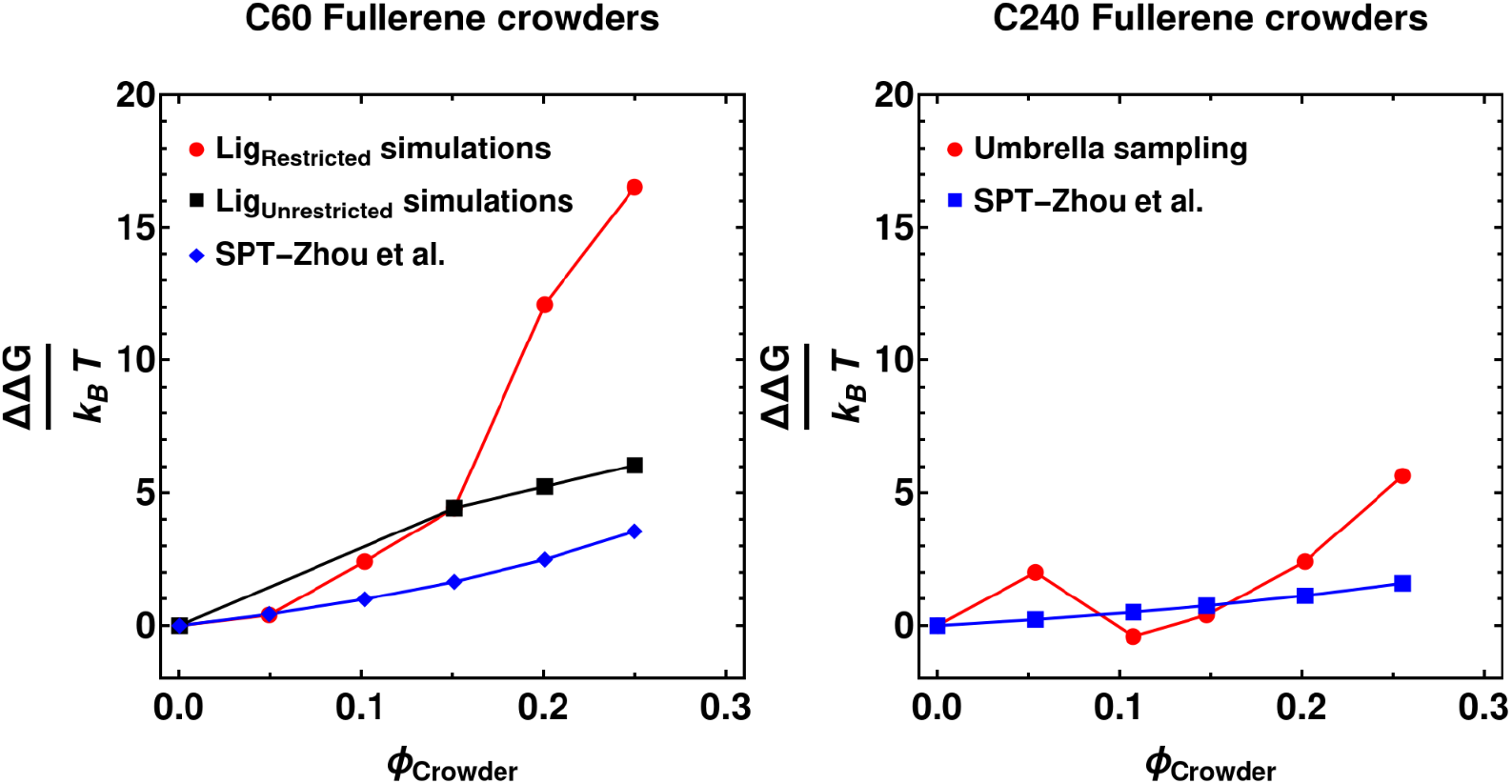
Comparison of ΔΔ*G* (k_B_T) of ligand-binding process in presence of C60 (panel (A)) and C240-crowders (panel (B)) at various crowder volume fraction in relative to dilute solution with SPT predictions. For C60-crowding we have also shown the data for unrestricted ligand movement cases for *ϕ*_*Crowder*_ = 0.15, 0.2 and 0.25.

For smaller crowder i.e. C60-fullerene, SPT predictions quantitatively agree with simulated values at very low volume fractions. However, with increase in crowder volume fraction, simulated ΔΔ*G* values become much larger compared to SPT predicted values. In contrast to C60, for larger crowder, C240, the quantitative agreement between simulated and SPT predicted ΔΔ*G* values are relatively better. Thus, it has been shown that for this particular ligand-binding process, smaller crowders can impose a much stronger effect compared to a larger crowder. Our simulation results for both the crowders are able to qualitatively reproduce the SPT predictions. In case of smaller crowders SPT largely underestimates results obtained from simulations. The quantitative disagreement between simulations and SPT can be referred to multiple reasons: (1) solvent contribution is not taken into account in SPT, (2) SPT prediction for ligand-binding equilibrium only depends on crowder-ligand interactions and relative sizes of crowder and ligand and (3) all components (crowder, ligand) are assumed to be hard spherical/spherocylindrical particles. In reality, ligand molecules do not act as inert hard particles. In our simulations ligand weakly interacts with the cavity atoms instead of being completely inert. One could argue that such effect must cancel out when we consider ΔΔ*G* as we subtract free energy contributions in crowded solution from that of in dilute solutions. However, we have successfully shown that even in presence of weak attractive interactions between ligand and cavity atoms, binding process becomes much more facilitated in crowded solutions compared to dilute solutions.

Role of water molecules is of utmost importance in bimolecular processes. We have explored the role of water molecules trapped inside the cavity (pocket water) on ligand-binding process. Earlier the role of water molecules have been extensively investigated in similar host-guest recognition processes in dilute solutions for sterically constrained ligand movement.^26–28^ It has been shown that unbinding of the ligand from the receptor site is accompanied by increase in number of pocket water molecules after the ligand reaches a critical distance from the receptor. Pocket water molecules (*n*_*w*_) are defined as the water molecules that are present inside the spherocylindrical pocket (please see Fig. 2). Here, we have investigated how such solvent assisted binding/unbinding process is further influenced by addition of large number of C60-Fullerene crowders. Mondal et al.^27^ reported large fluctuation in pocket water number in addition to slow relaxation of pocket-water fluctuation time-correlation function at a critical cavity-ligand distance where the ligand starts to transition from bound to unbound state. In order to gain a better understanding of the role of cavity-water molecules, we have calculated a two-dimensional free energy surface from the joint probability distribution of *n*_*w*_ and receptor-ligand distance from umbrella sampling simulations for restricted ligand movement. We have calculated the joint probability distribution from umbrella sampling trajectories as described in Ref. 50,51. The data are shown in supplementary information (Fig.S3)

In Fig. S3 we have shown two dimensional free energy surfaces as function of two reaction coordinates: number of pocket-water molecules and receptor-ligand distances obtained from umbrella sampling simulation trajectories for dilute solution and one of the crowding simulations. Large fluctuation in number of pocket-water molecules can be observed in dilute solution at approximately around receptor-ligand separation distances of 1.25-1.35 nm and 1.4-1.5 nm respectively. However, such pocket-water fluctuation is absent in presence of crowding. This shows that while the ligand binding process is associated with drying of the semi-ellipsoidal cavity both in presence and absence of crowding, in crowded environment (*ϕ*_*Crowder*_ = 0.25), the cavity remains mostly dry or contains very few water molecules on average, precluding the possibility of large water fluctuations and hence a wet pocket. Most noticeable difference between the two free energy profiles is observed for configurations at large cavity-ligand distances. For crowded condition, those states containing large number of pocket-water molecules (*n*_*w*_ *>* 16) are much higher in free-energy compared to that of the dilute solution (*ϕ*_*Crowder*_ = 0.0) case. This can be attributed to the fact that in crowded solution even when the ligand is at a large distance from the cavity, path of incoming water molecules towards the pocket is more or less obstructed by crowders. One can assume that those incoming water molecules may enter the hydration shell of the crowder molecules and become inaccessible to the semi-ellipsoidal pocket region. In addition to this, diffusion of water molecules in the system is largely going to be impeded due to high crowding concentration.^52^ A combined effect of these factors result in stabilization of dry configurations or configurations with fewer pocket-water molecules. Such predominant occurrence of a dry cavity or stability of states with small pocket-water molecules is a clear indication of the fact that ligand binding process is much strongly facilitated in presence of crowding compared to a dilute solution situation.

### Impact of crowding on freely diffusing guest ligand towards host

In the previous sections we have demonstrated the impact of macromolecular crowding on ligand-binding process involving a sterically restricted ligand. Here, we explore the effect of crowding on a similar process involving a freely diffusive C60-ligand molecule. For this purpose we have performed multiple unbiased and biased MD simulations allowing the ligand to move freely in any direction towards or away from the host. Please see the methods section for detailed simulation protocols.

Fig. 9, depicts the time evolution of cavity-ligand distances in presence and absence of C60-sized crowders when the ligand is freely diffusing in solution. In contrast to restricted ligand movement case, here we have obtained multiple binding events in dilute solution (*ϕ*_*Crowder*_=0.0) starting from a very large cavity-ligand separation of approximately 3.6 nm. In fact, a first-hand comparison of host-guest recognition trajectories (Fig. 9) between dilute (*ϕ*_*Crowder*_=0.0) and a relatively crowded (*ϕ*_*Crowder*_=0.25) does not reflect qualitatively significant acceleration on kinetics of binding time (so called *on-rate*). Accordingly, for a more rigorous and quantitative analysis of binding equilibrium, we have further performed two-dimensional umbrella sampling simulations (see methods for details) to map the free energetics of ligand movement in dilute and crowded conditions. Taking cue from the previous investigation,^26^ we have chosen receptor-ligand distances in the *z*-direction and in the *xy* plane (*ρ*) as reaction coordinates to describe ligand-binding process.

**Figure 9:**
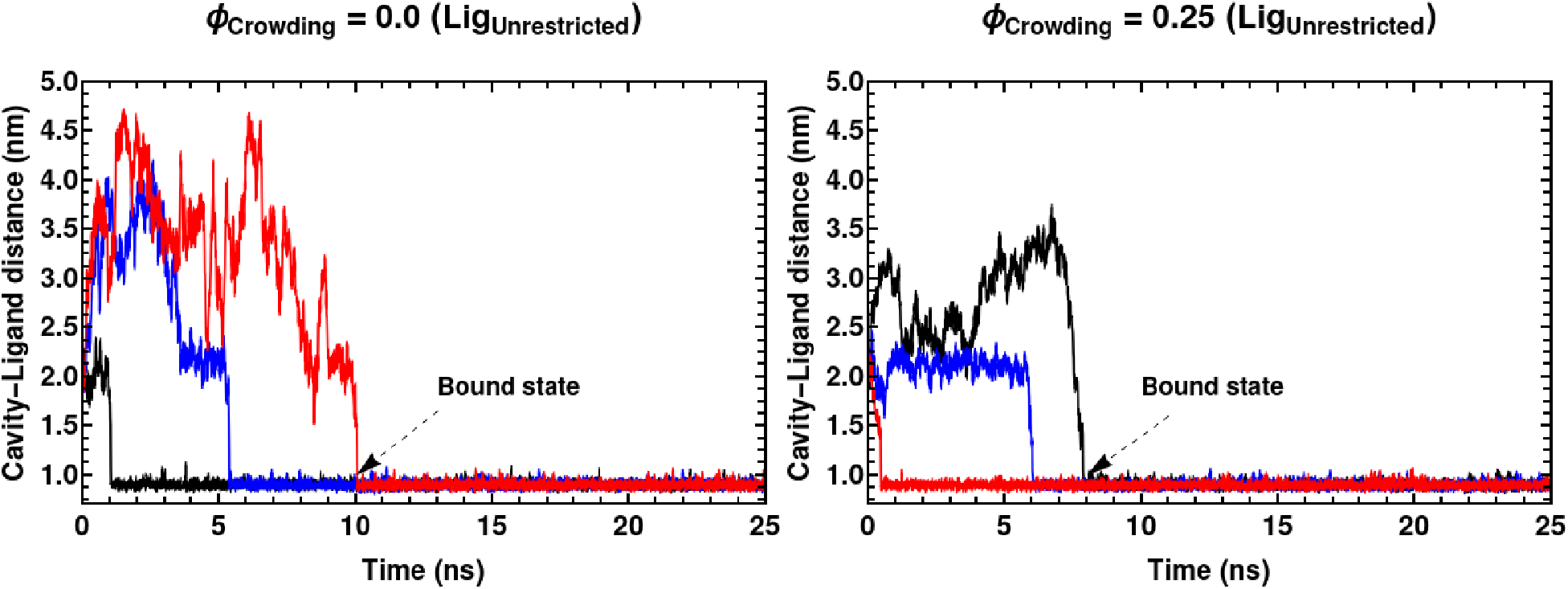
Time evolution of cavity-ligand distances obtained from unbiased simulations of freely diffusing ligand in presence of C60-fullerene crowder volume fractions (*ϕ*_*Crowder*_) = 0 and 0.25; Multiple independent unbiased MD simulation trajectories are shown together (red, black and blue lines), dashed arrows indicate appearance of bound states in the time profile.

In Fig. 10 we have shown the two-dimensional free energy surfaces obtained while the guest is free to move in any direction around the host receptor. A *ρ* value of zero indicates centrosymmetric approach of the ligand similar to restricted ligand movement case. Any non-zero value of *ρ* will refer to deviation from centrosymmetric approach of the ligand. The free energy minima in Fig. 10 are located at ∼ *z* = 0.83 nm and ∼ *ρ* = 0.35 nm which is slightly off-center. In Fig. 10 (inset) multiple unbiased MD trajectories were projected on the free energy surface. The bound, transition state(s) and unbound state(s) are shown in panel (B) of Fig. 10. The projected trajectory data approximately refer to a pathway (shown by dashed arrows in the Fig. 10 (inset)) where the ligand slides along the sides of the cavity and rolls in and out of the cavity during the binding-unbinding process. Centrosymmetric ligand-binding pathway is energetically unfavourable. The ligand follows a slide-and-roll approach and sidewise pathway^26^ for binding.

**Figure 10:**
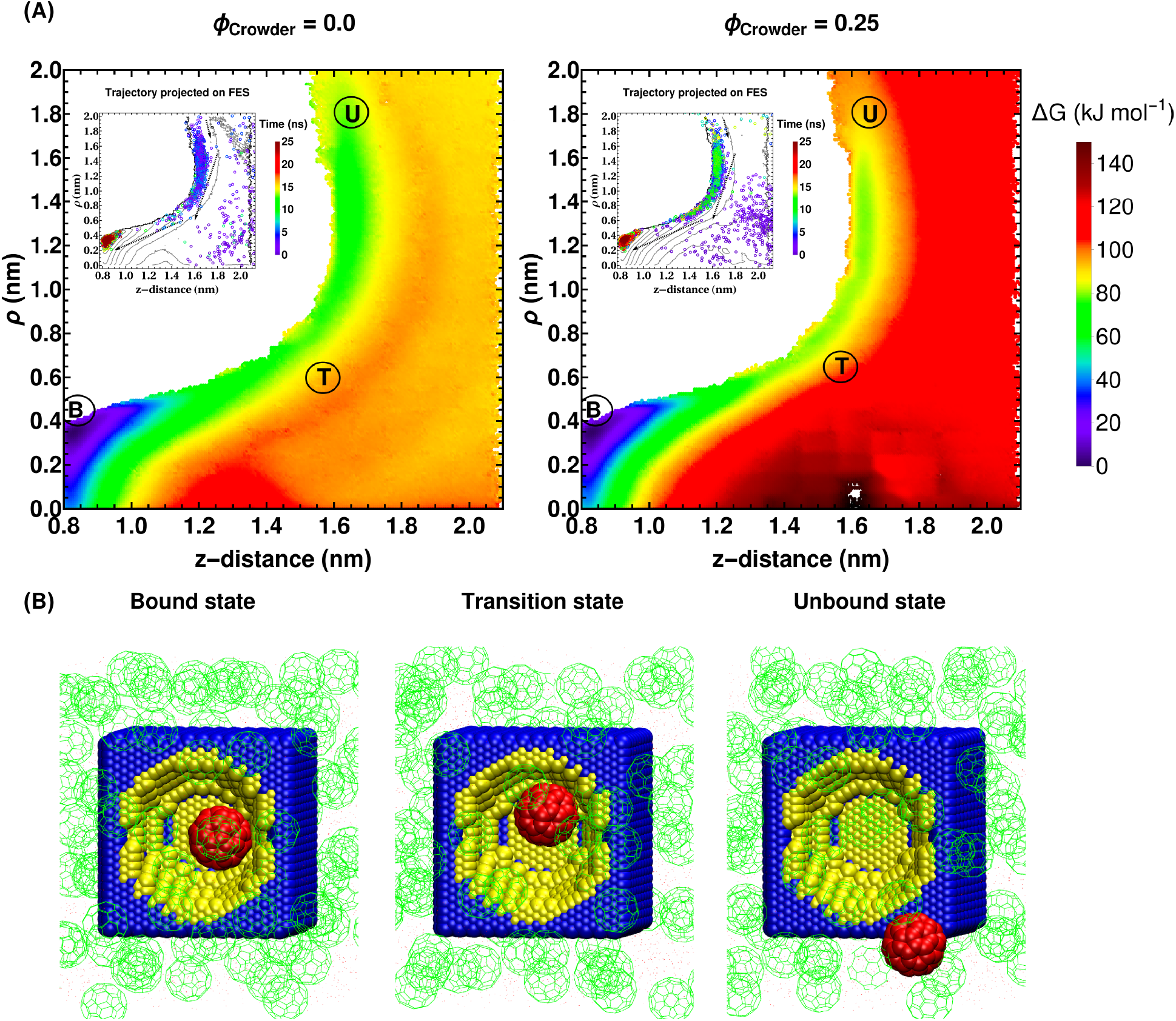
2D-free energy surface obtained from umbrella sampling simulations in (A) absence (*ϕ*_*Crowder*_ = 0.0) and (B) presence of crowding (*ϕ*_*Crowder*_ = 0.25), where the ligand movement is unrestricted. The labels **B, T and U** refer to the bound, intermediate and unbound states. Multiple independent unbiased MD simulation trajectories for *ϕ*_*Crowder*_ = 0 and 0.25 were projected onto the free energy surface; the projected data are colored by simulation time (ns)

In order to explain the ligand binding pathway, we have calculated the LJ interaction energies between the receptor and the ligand. We have shown a 2D-potential energy surface of ligand-receptor interactions in Fig. S5, which clearly depicts that the minimum energy pathway for the host-guest recognition process is along the sides of the receptor contrary to a centrosymmetric pathway. This can be attributed to the fact that the ligand is able to maintain stronger and distant hydrophobic interactions when approaching the receptor sidewise even after unbinding to a certain degree as shown in Fig. S5. The projection of the kinetic trajectories on the free energy surfaces (Fig. 10, inset) along with two approximately similar potential energy surfaces in panel (A) and (B) of Fig. S5, clearly shows that the ligand binding pathway remains more or less conserved in presence and absence of crowders. However, we also note that in complex biological environments in presence of many different interactions in addition to hard-core steric repulsion, the major ligand-binding pathways can get altered in presence of crowding, as has been demonstrated recently.^23^

As would be expected from excluded-volume driven crowding effect, the value of Δ*G*_*binding*_ or free energy of binding (≃ 100 kJmol^−1^) in case of unrestricted ligand movement, is also relatively larger in presence of crowders compared to dilute solution situation (≃ 85 kJmol^−1^). For unrestricted ligand movement, simulated ΔΔ*G* values (see Fig. 8) show much better quantitative agreement with SPT predictions compared to that of restricted ligand movement simulation results.

We have investigated the profile of cavity-water for the unrestricted ligand movement cases. The data are shown in the supplementary information (Fig.S4). For unrestricted ligand movement, we have found that in contrast to restricted ligand movement case the sharp cavity-water fluctuation is absent. Rather the ligand binding process is assisted by a gradual decrease of cavity-water. However, in this case as well, the crowding-induced enhancement of binding affinity is accompanied by drying of pocket.

A close comparison of crowding-induced change in binding free energy,(Table 3) across two scenarios (restricted versus unrestricted ligand movement) investigated here, clearly predicts that the effect of crowding would be significantly low when the ligand is unrestricted to move around the cavity, compared to the case when the ligand is sterically required to move along a centrosymmetric direction: the percentage increase in free energy of stabilisation is ∼ 18 % in unrestricted case versus ∼ 68 % in restricted case. For a focussed investigation to the possible origin of the significantly different crowding effect due to difference in ligand-movement, we opted for a kinetic dissection of the recognition process for largest crowder concentration and dilute solution (*ϕ*_*Crowder*_ = 0 and 0.25).

**Table 3:**
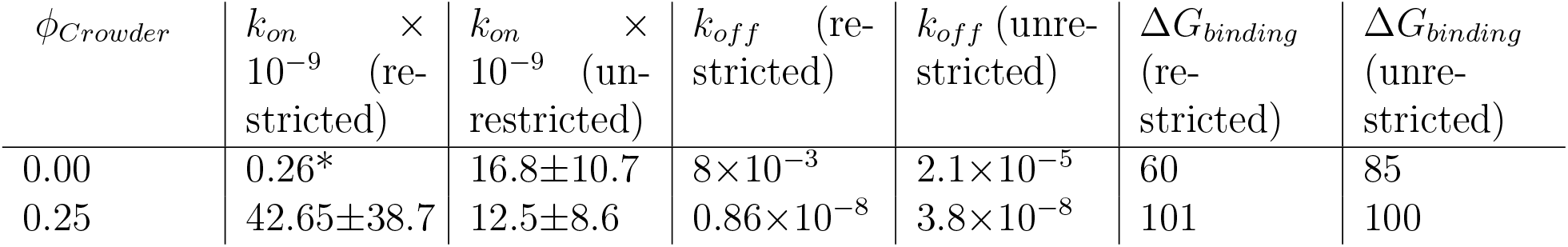
Crowder volume fractions (*ϕ*_*Crowder*_), *k*_*on*_ (M^−1^s^−1^), *k*_*off*_ (s^−1^) and Δ*G*_*binding*_ (kJ mol^−1^) obtained for restricted and unrestricted ligand movement. *values obtained from Ref. 28

### A kinetic analysis sheds light on impact of crowding on host-guest complexation

In our quest for the traits of significantly lesser impact of crowders in unrestricted ligand binding case, we have extracted the kinetic rate constants from our simulation results. In particular we computed binding rate constants i.e. on-rate constants (*k*_*on*_) and unbinding rate constants i.e. off-rate constants (*k*_*off*_). The ratio of these two rate constants is connected to Δ*G*_*binding*_ via a logarithmic function (see method),

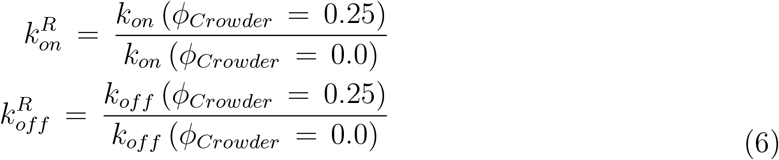

We find that in case of restricted movement (Table. 3), while the ligand binding on-rate constant (*k*_*on*_) increases, the unbinding rate constant (*k*_*off*_) decreases in presence of inert crowders, the extent of slow-down of off-rate constants is approximately ∼10^6^ times higher in magnitude relative to kinetics in dilute solution. On the other hand, when the guest ligand is free to move in any direction, there is almost no change or slight decrease in *k*_*on*_ due to presence of crowders. This is in line with the observation that the barrier to binding or the free energy cost of transitioning from the unbound state to the transition state (≃ 13-17 kJ mol^−1^) does not vary significantly for both the crowded and dilute solutions in case_*off*_ of unrestricted movement of the ligands. On the other hand, the observed crowding-induced free-energetic stabilization of the binding event during unrestricted movement of ligand is mostly driven by decrease in *k*_*off*_. However, the extent of slow-down in *k*_*off*_ is significantly lower during unrestricted ligand movement than the restricted ligand movement, thereby resulting in less muted impact of crowders in binding affinity. In Table. 4, we have shown the relative on and off rate constants for restricted and unrestricted ligand movement. Here, the relative rate constants: 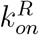 and 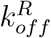 are defined as the ratio of either on-rate or off-rate constants in presence and absence of crowders as described in Eqn. 6. It can be seen from Table. 4 that the extent of relative decrease in off-rate constant is much stronger than the increase in on-rate constant in crowded environment when compared to dilute solution. It is clearly evident from Table. 4 that the impact of steric restriction on ligand movement is much more stronger on the off-rate constant than the on-rate constant in presence of crowding when compared to dilute solution rate constants. Together these results suggest that crowders impact the ligand residence time (which is the inverse of *k*_*off*_) the most, rather than accelerating the binding process. Most importantly, the impact in ligand residence time is considerably less in case of unrestricted ligand movement.

**Table 4:** Relative on 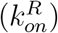 and off 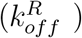 rate constants obtained for restricted and unrestricted ligand movement; upward and downward arrows refer to the increase or decrease of the on or off rate constants in crowded environment with respect to dilute solution

| Ligand movement | $k_{on}^R$ | $k_{off}^R$ |
| --- | --- | --- |
| Restricted | 164.04 (↑) | $1.075 \times 10^{-6}$ (↓) |
| Unrestricted | 0.74 (↓) | $1.81 \times 10^{-3}$ (↓) |

A possible justification of this observation can be argued in terms of the configurational entropy of the ligand in both situations. Ligand binding process is accompanied by reduction of entropy of the ligand. In presence of crowding, for sterically constrained ligand movement, the decrease in ligand entropy upon binding will be much less as the ligand entropy in the unbound state will be much smaller due to the a priori steric restrictions on ligand movement. However, for the unrestricted ligand movement case, in absence of any a priori steric restrictions on the ligand movement, the ligand entropy will be much larger at the unbound state. Thus, the ligand-binding process will be entropically more unfavorable when the ligand is freely diffusing in any direction. We propose that the entropic destabilization will be dominant over the enthalpic stabilization of the ligand-bound state resulting in weaker relative impact of crowding on ligand-binding process when the ligand is free to move in any direction.

## Summary

In this study we have demonstrated the impact of macromolecular crowding on ligand binding process of a previously studied prototypical substrate-ligand system. We have studied thermodynamic and kinetic aspects of a sterically constrained and freely diffusing ligand binding-unbinding process using unbiased and enhanced-sampling MD simulations in explicit water with full atomistic resolution. Introduction of repulsive macromolecular crowders are shown to increase the free energy of binding, barrier towards unbinding and reduce the barrier height towards binding via excluded volume effect. The facilitation of the binding is also reinforced via crowder-induced desolvation of the host receptor. Furthermore, we have studied the impact of different sizes of crowding agents on this molecular recognition process. It has been shown that smaller-sized crowders (C60-Fullerene crowder) influence the ligand binding equilibrium much strongly compared to larger crowders (C240-Fullerene crowder). This can also be argued in terms of more favorable or less unfavorable crowder-ligand interactions present for chemically soft repulsion in larger-sized crowders. However, when the ligand is free to move in any direction, the free energy calculations show that while free energy of binding increases in presence of crowding, the barrier to binding does not change significantly compared to dilute solution case. This clearly depicts that crowding does not significantly perturb the ligand-binding equilibrium with respect to dilute solution for freely diffusing ligand. It has also been shown that for this prototypical host-guest system, the ligand binding pathway is conserved in presence and absence of crowding. However, for complex biological systems the ligand binding pathway can be altered due to the presence of a multitude of intermolecular interactions apart from steric repulsion. ^23^ Furthermore the calculation of *k*_*on*_ and *k*_*off*_ shows that for both restricted and unrestricted ligand movement, the impact of increasing crowding intensity on *k*_*on*_ is either relatively small (in case of constricted guest movement) or almost muted (in case of unrestricted guest movement). In contrast, the key to crowding-induced stabilisation of host-guest complex lies in significant slow-down of *k*_*off*_ or increase in guest-residence times inside the host, compared to dilute solution. The concomitant increase in residence times inside the host due to the crowders is much larger when the guest is constricted to approach the host along a direction, than when freely diffusing.

We have further compared the values of change in free energy of binding or ΔΔ*G*_*binding*_ obtained from simulations with analytical predictions obtained using SPT of hard sphere fluids. Simulation results are shown to qualitatively agree well with the SPT-predictions. SPT-can be successfully employed for analytic description of crowding effect for crowders from freely diffusing ligand movement. However, for restricted ligand movement, SPT is shown to underestimate the simulated results strongly. One of the major features of our study is the inclusion of water molecules explicitly while simulating crowded environment, a feature that has often been overlooked in past. Ligand binding in this prototypical cavity-ligand system has been shown to occur with sharp dewetting of the binding pocket. We have also investigated the role of cavity-water molecules in ligand binding. The 2D-free energy profiles as a function of receptor-ligand separation and number of pocket water molecules show that semiellipsoidal pocket region predominantly remains dry or contains very few water molecules in presence of high C60-Fullerene crowder concentration. This effectively reinstates our conclusion that ligand binding is strongly facilitated in strongly crowded environments. As the ligand approaches the cavity, water molecules at the ligand-cavity interface gets depleted as a consequence of depletion effect as described in Asakura-Oosawa theory of depletion forces.^53^

In summary we have presented thermodynamic and kinetic insights of molecular recognition process for a well studied prototypical cavity-ligand system in presence and absence of macromolecular crowding using hard-spherical crowders in explicit solvent with full atomistic resolution. We have studied the impact of crowder size as well as the impact of sterical constraint to ligand movement. We propose that impact of macromolecular crowding will be much more stronger when ligand movement is largely restricted e.g. in cases of small molecule binding to and translocation through ion-channels, membrane proteins etc., specific and non-specific binding of transcription factors to chromosomes and DNA^40^, when compared to the unrestricted free movement of the ligand.

This simple model system can be used to make useful predictions on crowding effects on realistic molecular recognition processes. However, that would require further refinement of this model system. For example, the current study does not consider any enthalpic contribution to the overall crowding effect. It has been shown in many recent experimental and simulation studies that a non-specific attractive interaction between crowders and solute molecules can lead to very small changes to almost no change and mild stabilization of unfolded macromolecular configurations. ^7,49,54–57^ In future this simple model system can further be extended incorporating attractive crowder-ligand interactions making this a more realistic model system for predicting impact of macromolecular crowding on realistic protein-ligand binding processes.

## Supplementary Information

All supplemental figures described in the article (PDF)

## Acknowledgments

This work was supported by shared computing resources obtained from TIFR Hyderabad, India. We acknowledge support of the Department of Atomic Energy, Government of India, under Project Identification No. RTI 4007. JM acknowledges Core Research grants provided by the Department of Science and Technology (DST) of India (CRG/2019/001219). BBM acknowledges Satyabrata Bandyopadhyay and Mohammad Sahil for useful discussions.

