## Supplemental Figures for "Impact of Inert Crowders on Host-Guest Recognition Process"

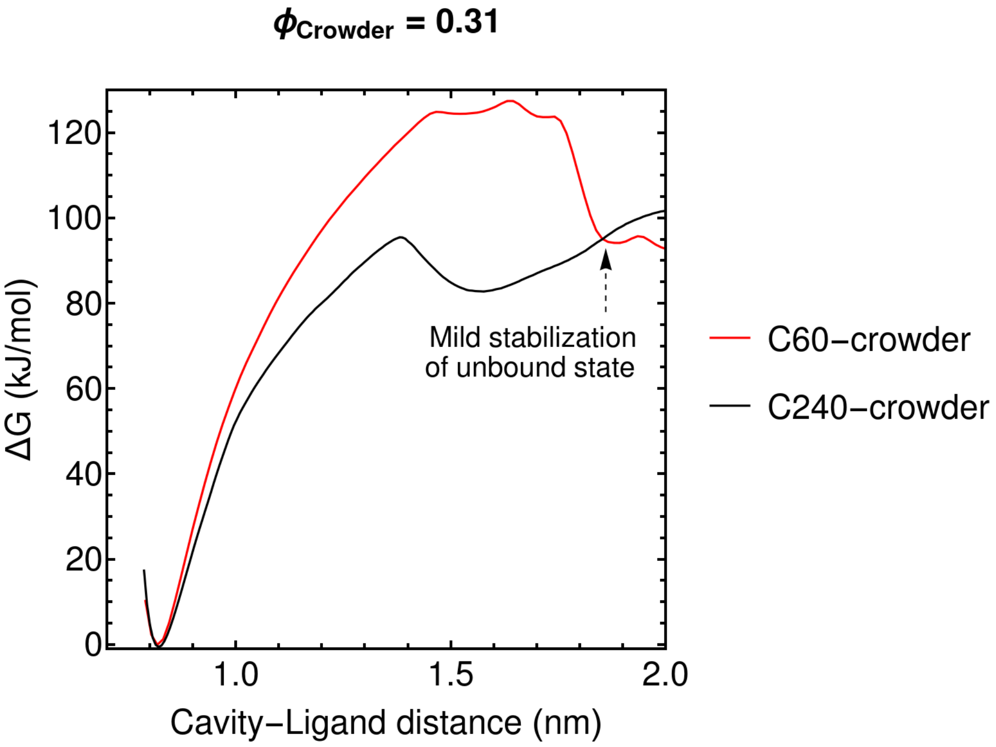


Figure S1: Free energy profiles as a function of cavity-ligand distances obtained for high crowding concentration, $\phi_{crowder} = 0.31$ for C60 and C240-fullerene crowders; dashed arrow refers to a mild stabilization of unbound states occurring in presence of C60-fullerene crowder

In Fig. S1, we have shown free energy profiles obtained from biased umbrella sampling simulations for $\phi_{crowder} = 0.31$ in presence of C60 and C240-fullerene crowders. For C60-

fullerene profile (Red line) a mild stabilization of the unbound states starts to appear at large ligand and receptor separations.

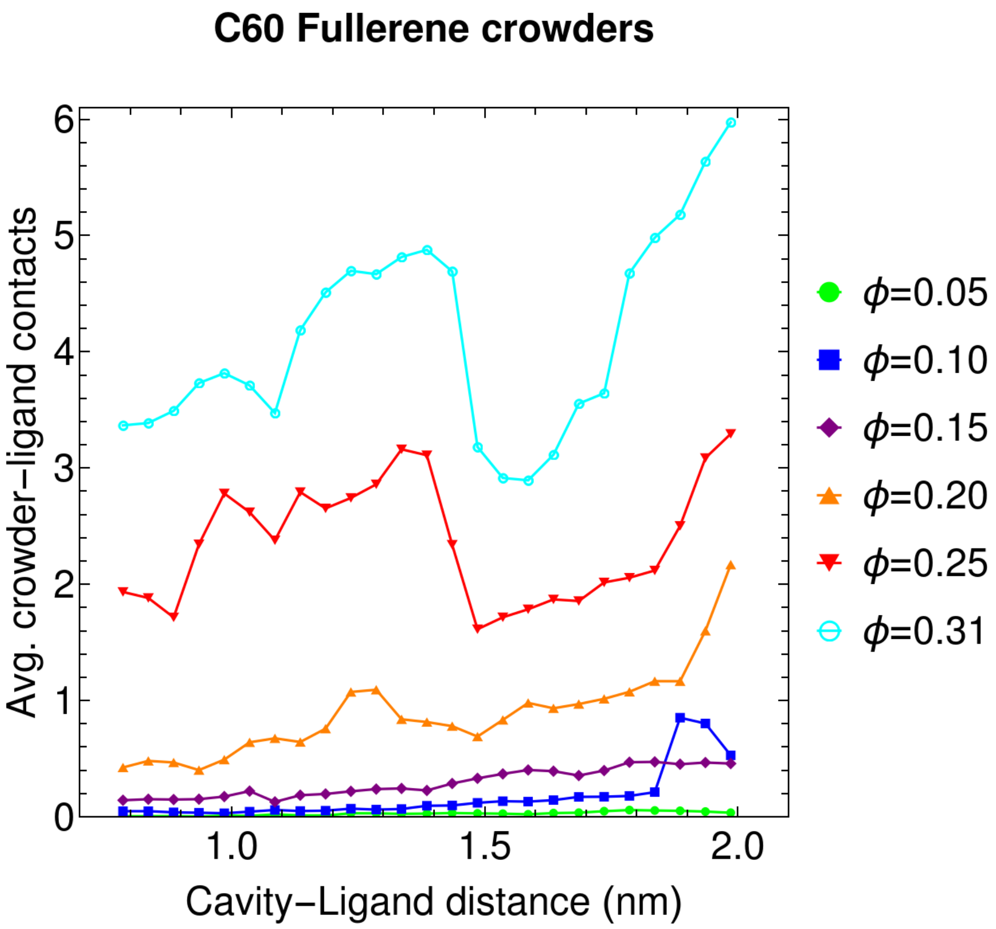


Figure S2: Average crowder-ligand contact obtained for various concentrations of C60-fullerene crowder at different receptor-ligand separations.

In Fig. S2, we have shown average number of crowder-ligand contacts obtained from biased umbrella sampling simulations for various C60-fullerene crowding concentrations where ligand movement is restricted. Large number of crowder-ligand contact starts to appear for high crowder concentrations ($> 0.2$) at large receptor-ligand separations ($> 1.8$ nm).

In Fig. S3, a 2D free energy surface is shown as a function of the number of pocket water molecules and receptor-ligand distances in crowded and dilute solutions for sterically restricted ligand movement. The host-guest recognition process is assisted by a sharp fluctuation in pocket-water number at ligand-receptor separations of 1.25-1.35 nm and 1.4-1.5

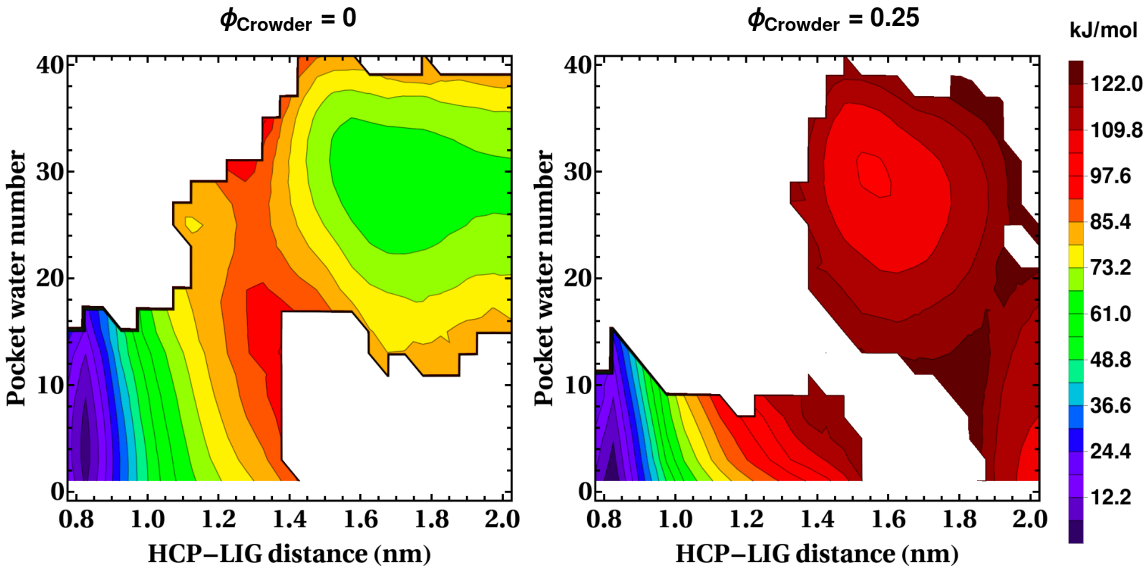


Figure S3: Two dimensional free energy surface as a function of number of pocket-water and receptor-ligand separation distances for dilute solution ($\phi_{crowder} = 0$ and in presence of C60-Fullerene crowders ($\phi_{crowder} = 0.25$)) for restricted ligand movement

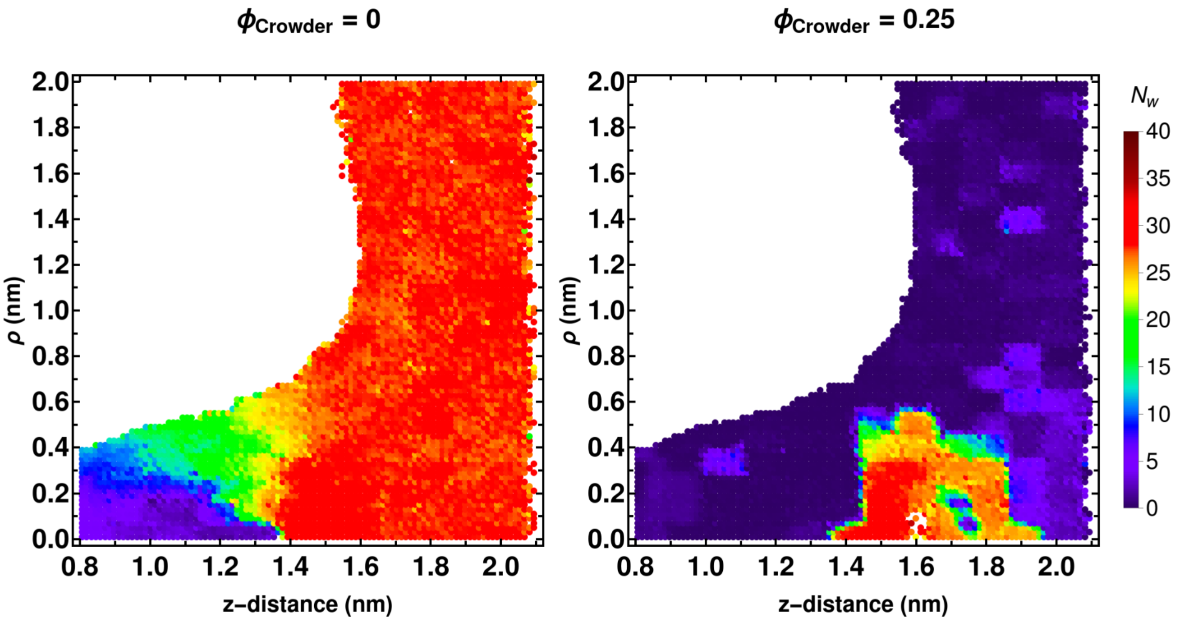


Figure S4: Two dimensional scatter plot of $\rho$ vs $z$ as a function of number of pocket-water molecules for dilute solution ($\phi_{crowder} = 0$ and in presence of C60-Fullerene crowders ($\phi_{crowder} = 0.25$)). The data points are colored based on $N_W$, pocket water number.

nm respectively. Crowding induced ligand binding is mostly facilitated by the drying of the host binding pocket resulting from obstruction of the pocket by large number of crowders.

In Fig. S4 we have shown the a 2D-scatter plot of reaction coordinates ($z$ and $\rho$) for unrestricted ligand movement in presence and absence of crowding, as a function of number of pocket-water molecules. Pocket-water or cavity-water molecules are described as the water molecules trapped inside the binding pocket. For detailed description please refer to the main text. Ligand binding in dilute solution (left panel) is assisted by a gradual dewetting of the cavity or gradual stripping of cavity-water molecules in contrast to restricted ligand movement case (assisted by sharp dewetting of the cavity) In crowded solution (right panel), cavity remains mostly dry indicating a predominant occurrence of ligand-bound states. There are some patches at $z$ < 1.4-1.8 nm and $\rho$ < 0.3 nm, which contains large number of cavity-water molecules. However, these states are very high in energy (refer to main text for 2D-Free energy surface) due to the near centrosymmetric approach of the ligand and are less statistically significant.

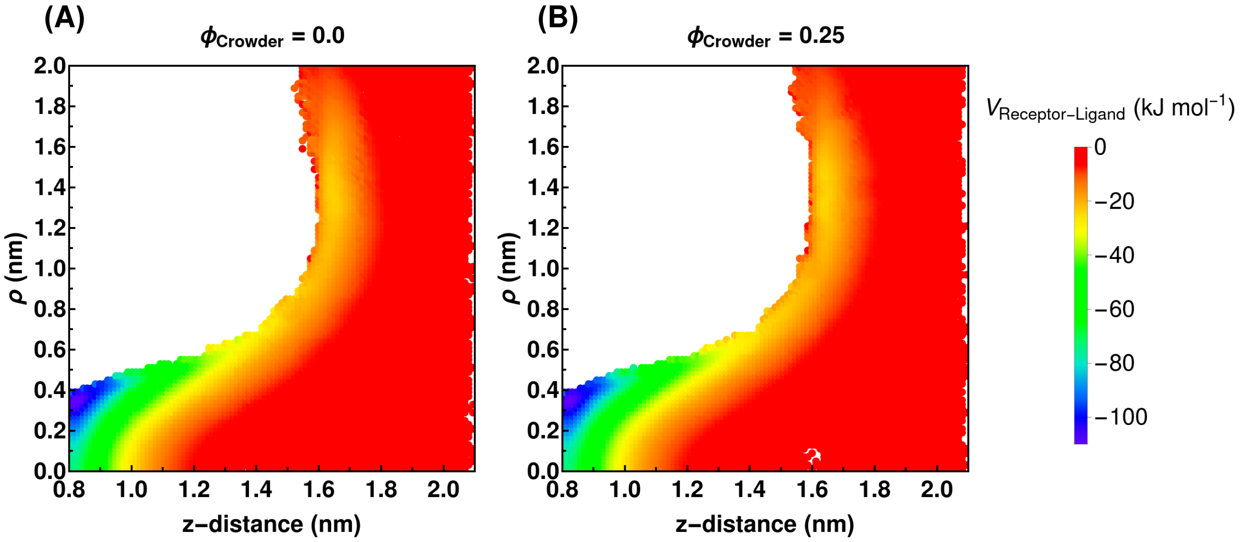


Figure S5: 2D-potential energy surface (receptor-ligand LJ interactions) obtained from umbrella sampling simulation trajectories in (A) absence ($\phi_{Crowder}$ = 0.0) and (B) presence of crowding ($\phi_{Crowder}$ = 0.25), for freely diffusing ligand.
